# Genome-Wide CRISPR-Cas9 Screening Identifies a Synergy between Hypomethylating Agents and SUMOylation Blockade in MDS/AML

**DOI:** 10.1101/2024.04.17.589858

**Authors:** Peter Truong, Sylvie Shen, Swapna Joshi, Md Imtiazul Islam, Ling Zhong, Mark J. Raftery, Ali Afrasiabi, Hamid Alinejad-Rokny, Mary Nguyen, Xiaoheng Zou, Golam Sarower Bhuyan, Chowdhury H. Sarowar, Elaheh S. Ghodousi, Olivia Stonehouse, Sara Mohamed, Cara E. Toscan, Patrick Connerty, Purvi M. Kakadia, Stefan K. Bohlander, Katharine A. Michie, Jonas Larsson, Richard B. Lock, Carl R. Walkley, Julie A. I. Thoms, Christopher J. Jolly, John E. Pimanda

**Author notes:** Senior authors. Correspondence; Dr Christopher Jolly, Dr John Pimanda.

## Abstract

Hypomethylating agents (HMAs) are frontline therapies effective at altering the natural course of Myelodysplastic Neoplasms (MDS) and Acute Myeloid Leukemia (AML).

However, acquired resistance and treatment failure are hallmarks of HMA therapy. To address this clinical need, we performed a genome-wide CRISPR-Cas9 screen in a human MDS-derived cell line, MDS-L, and identified TOPORS as a highly ranked loss-of-function target that synergizes with HMAs, reducing leukemic burden and improving survival in xenograft models. We demonstrate that the depletion of TOPORS mediates sensitivity to HMAs by predisposing leukemic blasts to an impaired DNA damage response (DDR) accompanied by an accumulation of SUMOylated DNMT1 in HMA-treated TOPORS-depleted cells. Importantly, the combination of HMAs with targeting of TOPORS did not functionally impair healthy hematopoiesis. While inhibitors of TOPORS are currently unavailable, we show that inhibition of protein SUMOylation (upstream of TOPORS functions) with TAK-981 partially phenocopies HMA-sensitivity and DDR impairment. Overall, our data suggest that the combination of HMAs with the inhibition of SUMOylation or TOPORS demonstrates a favourable therapeutic index and is a rational treatment framework for High-Risk MDS (HR-MDS) or AML.

## INTRODUCTION

The cytidine nucleoside analogues Azacitidine (AZA) and Decitabine (DAC) are effective frontline treatments for MDS which promote hematologic recovery and delay transformation to AML^1–4^. Despite their clinical benefit, HMA therapy is limited by transient efficacy as underscored by the high frequency of acquired resistance to recurrent HMA exposure^2^ and disease relapse. Emerging evidence suggests multiple mechanisms of HMA resistance; namely, adaptations of metabolic processes involved in activating HMAs^5^, cell cycle quiescence^6^, disequilibrium between pro- and anti-apoptotic proteins^7^, upregulation of immune checkpoint signalling axes^8^, and re-expression of oncogenes^9^. Allogeneic bone marrow transplants, the only potential curative approach, are feasible for approximately 8% of MDS patients due to their typical frailty^10^. Current treatment options for non-responding patients are limited to enrolment into clinical trials or provision of supportive care.

Strategies to develop combinatorial treatments to overcome acquired drug resistance and tumor heterogeneity have been effective across various cancer types^11^. In MDS and AML, combining the BCL-2 inhibitor Venetoclax (VEN) with AZA is effective in eradicating malignant leukemic stem cells compared to monotherapy^12^. However, the associated extreme rates of febrile neutropenia are of high concern^13,14^. Combining anti-CD47 (magrolimab) with AZA has also been explored. Preliminary clinical data suggested promising survival benefits, especially in *TP53*-mutated subsets^15^, but Phase 3 ENHANCE studies combining magrolimab with AZA (NCT04313881) in HR-MDS, and magrolimab with VEN and AZA in AML (NCT05079230) were discontinued due to futility and increased mortality compared with AZA or AZA and VEN, respectively.

Current models of drug mode-of-action for HMAs include rewiring of the epigenome to promote the re-expression of silenced tumor suppressor and cellular differentiation genes^16^, the induction of endogenous retroviral elements leading to inflammatory viral mimicry responses^17,18^, and cytotoxicity through the formation of genotoxic covalent DNMT1-DNA adducts^19^. To rationally identify secondary agents for combinatorial therapy, the epistatic genetic interactions that define drug response for the anchoring agent need to be systematically mapped. Genome-wide screening approaches can identify synthetic lethal relationships in an unbiased manner without requiring prior knowledge of drug mechanism. Although genome-wide CRISPR-Cas9 screens^20,21^ and targeted RNAi screens to identify resistance and synthetic lethal HMA-gene relationships^22^ have been performed, a genome-wide CRISPR-Cas9 dropout approach to identify genetic vulnerabilities in MDS cells to low dose HMA therapy has not yet been reported.

We performed a genome-wide loss-of-function CRISPR-Cas9 dropout screen in *TP53* mutant MDS-L cells in the presence of low-dose AZA and identified the E3-ligase TOPORS as a top sensitization target. In the absence of an available TOPORS inhibitor, we provide genetic proof of concept that targeting TOPORS confers hypersensitivity to HMAs by delaying the clearance of SUMOylated DNMT1 from HMA-treated cells, predisposing leukemia cells to apoptosis. Importantly, this strategy did not impair healthy hematopoiesis. As a surrogate for directly targeting TOPORS, we show that inhibition of SUMOylation using TAK-981 is synergistic with HMAs and phenocopies the DDR impairment observed in *TOPORS*-edited MDS cells without disrupting healthy hematopoietic stem and progenitor cells (HSPC) function. Our work reveals a unique therapeutic approach to enhance HMA response through modulating SUMOylation-dependent DDR, and more broadly, provides a framework for development of effective HMA combinatorial therapies.

## RESULTS

### Genome-wide CRISPR-Cas9 dropout screening identifies novel genetic determinants of AZA sensitivity

MDS-L was selected for genome-wide CRISPR-Cas9 dropout screen as it is the only MDS cell line that faithfully recapitulates MDS pathogenicity *in vivo*^23^ and harbors a genetic profile reflective of HR-MDS (del5q, *TP53*^-/-^, Type II myeloid driver mutations^24^, Table S1). We chose 0.3μM AZA for dropout screening because compared to higher concentrations it mediated anti-proliferative rather than direct lethality, while maintaining robust levels of demethylation (Fig. S1A). Cas9-expressing MDS-L cells were transduced with the Brunello sgRNA library^25^ and treated with AZA or vehicle for 12 cellular divisions in the AZA-treated arm (18 divisions in the vehicle-treated arm, Fig. S1B) to ensure robust depletion or enrichment of sgRNAs (Fig. 1A). The quality of the CRISPR-Cas9 screening libraries and the frequency of individual sgRNAs in the two treatment arms were quantified using the MAGeCKFlute analysis pipeline (Fig. 1B, Table S2)^26^.

**Figure 1.**
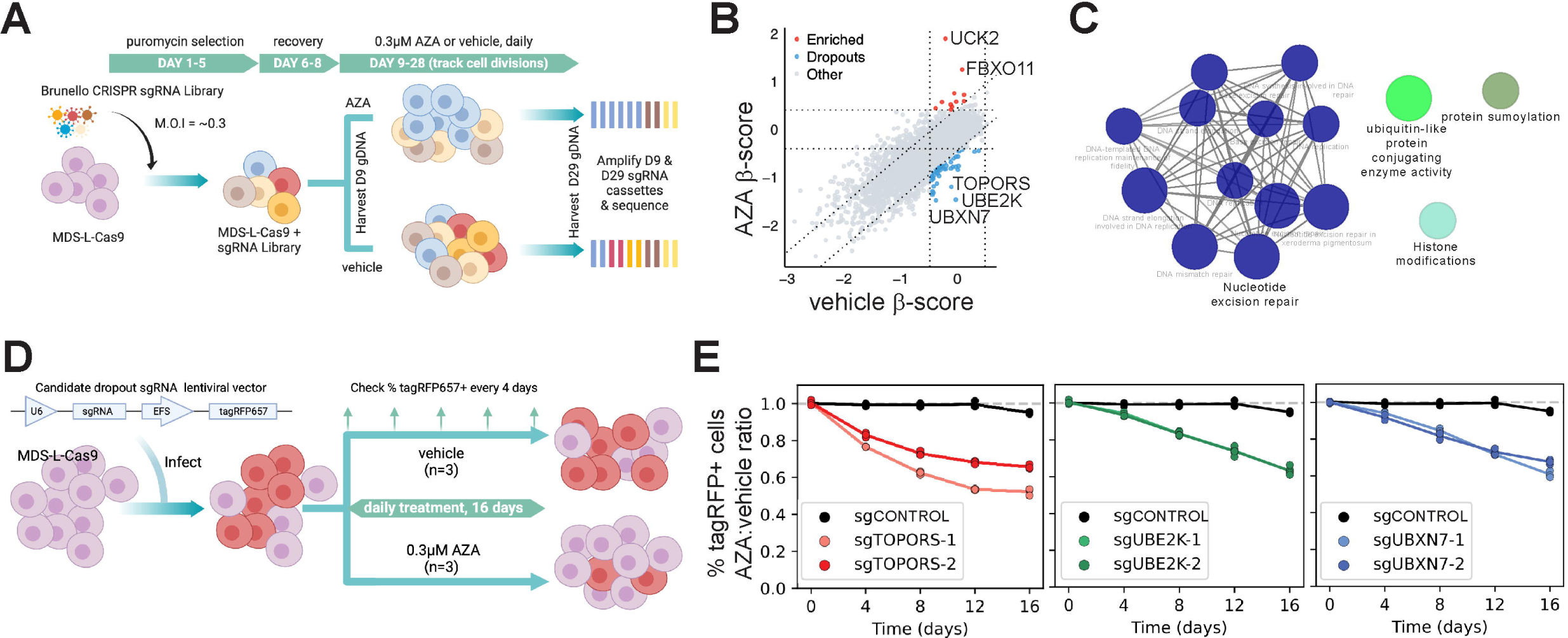
Genome-wide CRISPR-Cas9 dropout screening identifies novel genetic determinants of AZA-sensitivity. (A) Schematic of the genome-wide CRISPR-Cas9 dropout screen workflow performed in AZA-treated Cas9-expressing MDS-L. Image made with BioRender. (B) MAGeCKFlute nine-square correlation plot using cell-cycle normalized β scores calculated for each gene target (n=2). Colors: sgRNAs specifically (●) enriched, or (●) depleted under AZA selection. (C) ClueGO pathway term network highlighting biological processes enriched in dropout hits. (D) Competitive proliferation assay workflow. MDS-L/Cas9 cells were transduced with lentiviral vectors encoding a single sgRNA plus a tagRFP657 (tagRFP657) reporter from separate promoters. Image made with BioRender. (E) Validation of AZA-selection against *TOPORS-*, *UBE2K-*, and *UBXN7-*editing using the competitive proliferation assay shown in D. In each plot, y = %tagRFP^+^ in AZA/mean %tagRFP^+^ in vehicle; n=3.

As expected, we identified editing of *UCK2*^20,22^, which encodes the kinase that converts AZA from nucleoside to nucleotide, as the top hit conferring resistance to AZA. We also identified genes belonging to pathways involved in the DDR (*DYNLL1, ATMIN, BAX*), tumor suppression (*FBXO11, GFI1B*), mRNA processing (*PSIP1, SMG9, DDX3X*), and histone modification (*HDAC2, USP22*) as enriched hits. Fifty dropout gene targets conferring hypersensitivity to AZA were identified. We focused on the sgRNAs depleted in the AZA, but not vehicle, arm of our screen, as these their gene targets encode potential combination drug ligands. Gene ontology terms, KEGG pathways, and WikiPathways clustered dropout hits into key biological processes including excision DNA repair pathways, protein SUMOylation, histone modification, and ubiquitin-like protein conjugating activities (Fig. 1C). This was distinct from pathways significantly depleted in both AZA- and vehicle-treated arms of the screen (Fig. S1C). AZA-dropout hits were ranked according to their degree of depletion, revealing E2- and E3-ubiquitin ligases UBE2K and TOPORS, and the ubiquitin-interacting protein UBXN7 as top dropout hits (Fig. S1D). To validate the impact of individual gene perturbations on AZA sensitivity, two separate sgRNAs targeting each of *UBXN7*, *UBE2K*, or *TOPORS* were individually transduced into MDS-L-Cas9. ICE algorithm^27^ analysis of Sanger sequencing across expected Cas9-cut sites revealed high frequency polyclonal indels in the target genes with useful KO scores (Fig. S1E). Cellular proliferation was assessed by tracking the frequency of tagRFP657-expressing cells over time (Fig. 1D). In close concordance with our whole-genome screen, targeting *UBXN7, UBE2K*, or *TOPORS* resulted significant depletion of tagRFP657+ cells under AZA selection, validating these genes as *bona fide* drug target candidates that synergize with AZA therapy (Fig. 1E).

### Loss of TOPORS sensitizes leukemia cell lines to HMAs

Our top-ranked candidate was TOPORS. As an E3-ligase acting downstream of E1- and E2-ligases in ubiquitylation and SUMOylation pathways^28–31^, specific inhibitors of TOPORS could be expected to have fewer undesirable side effects than inhibitors of E1- or E2-ligases, making it the most attractive target from a theoretical therapeutic index standpoint. To test for specific relevance to blood malignancies, gene expression was queried using the TCGA database, which revealed that *TOPORS* is more highly expressed in human leukemia compared to other cancer types (Fig. S2A).

We next determined the AZA dose-response relationship using *TOPORS*-edited MDS-L cells across a range of AZA concentrations. Editing of *TOPORS* with two independent sgRNAs (Fig. S1E) sensitized MDS-L-Cas9 cells to four consecutive days of AZA treatment by up to 3.4-fold (Fig. 2A). For orthogonal validation, we generated MDS-L lines lentivirally expressing shRNAs against *TOPORS* (Fig. S2B), which showed a similar increase in sensitivity to AZA (Fig. 2B). Furthermore, enhanced sensitivity of *TOPORS*-edited cells to AZA therapy was associated with a marked synergistic reduction in clonogenicity (Fig. 2C).

**Figure 2.**
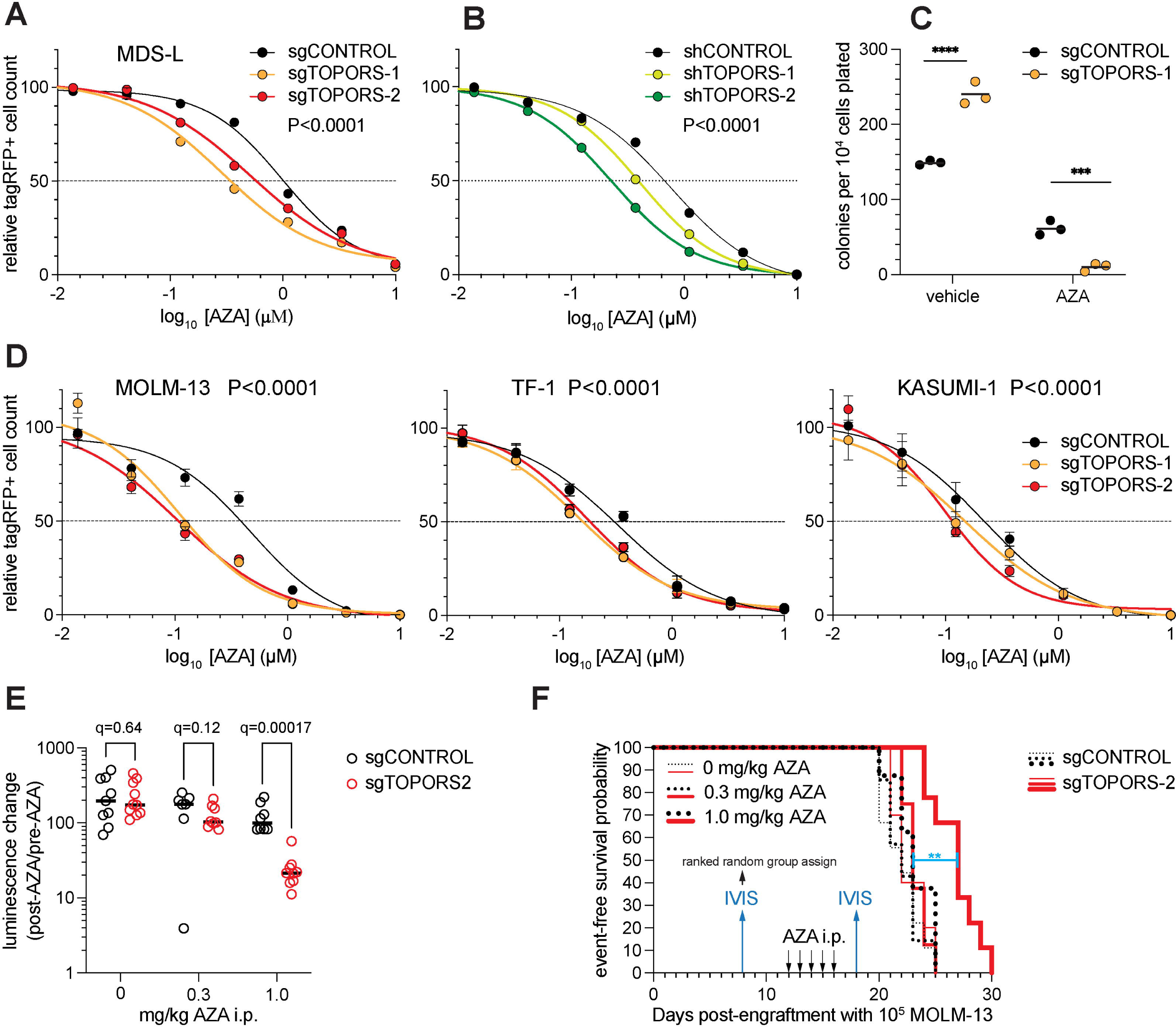
Loss of TOPORS sensitizes MDS and AML cell lines to AZA. (A-B) Dose-survival plots of tagRFP^+^ cell counts following four daily applications of AZA MDS-L cells polyclonally expressing Cas9 plus (A) single sgRNAs, or (B) single shRNAs which targeted *TOPORS* or a non-targeting control. Dots are means (n=4) normalized to the vehicle control, ±SD. P-value is from an extra sum-of-squares F test. (C) Clonogenic assays performed using *TOPORS*-edited MDS-L cells pre-treated with 0.3µM AZA as in A before plating in methylcellulose medium. Colonies were counted two weeks after methylcellulose plating. 2-way ANOVA: *** P≤ 0.001, **** P ≤ 0.0001. (D) Dose-survival plots of tagRFP^+^ cell counts following 4 days of daily treatment with the indicated AZA concentrations in AML cell lines polyclonally expressing Cas9 plus single sgRNAs or a non-targeting control sgRNA. Dots are means (n=4) normalized to the vehicle control, ±SD. P-values are from extra sum-of-squares F tests. (E) The change in whole body luminescence flux in MISTRG mice engrafted with 10^5^ MOLM-13 cells which polyclonally express luciferase, Cas9 and the indicated sgRNAs, immediately following 1 cycle of treatment with AZA or vehicle i.p.-as described by the time-based x-axis in F. FDR q-values (threshold = 0.01) are reported for a Mann-Whitney multiple comparison test; n=7–10. (F) Kaplan-Meier plots for survival of the same MISTRG mice as E. Whole body luminescence (“IVIS”) was performed 8 days after engraftment to give a “pre-AZA” baseline (which was used to rank-randomize mice into treatment groups based on sex and relative engraftment-“pre-AZA” in E), and again on day 18 – two days after completion of the treatment cycle (“post-AZA” in E). Event-free survival was scored according to ethics guidelines. ** Mantel-Cox test P=0.0029 for sgCONTROL versus sg*TOPORS-2* at 1.0mg/kg AZA.

As HMA therapy is used to treat a range of myeloid malignancies, we determined the applicability of targeting TOPORS in AML. We generated three *TOPORS*-edited AML cell lines (Fig. S2C) which reflect different stages of leukemic myeloid differentiation, including MOLM-13 (*TP53*^WT^ AML line derived from a patient with antecedent HR-MDS), TF-1 (*TP53*^WT^ erythroleukemia line with del5q aberrations), and Kasumi-1 (*TP53*^-/-^ mutant pediatric acute myeloblastic leukemia line). Polyclonal *TOPORS*-editing resulted in significant increases in AZA sensitivity in all cell lines (Fig. 2D). Furthermore, enhanced HMA-sensitivity conferred by *TOPORS*-editing extended to the 2’-deoxy derivative of AZA, Decitabine (DAC) (Fig. S2D), implying that TOPORS acts downstream of drug incorporation into DNA.

To evaluate the *in vivo* significance of our findings, we transduced *TOPORS*-edited and control MOLM-13 cells with lentivirus encoding luciferase-GFP^32^, then transplanted GFP^+^ sorted cells into cytokine-humanized adult “MISTRG” immune-compromised mice^33^, treated the mice for five consecutive days with AZA, and monitored disease progression through bioluminescence imaging and event-free survival. In recipients transplanted with *TOPORS*-edited MOLM-13 cells, a single five-day cycle of 1mg/kg i.p. AZA treatment transiently reduced leukemia burden relative to controls by about 90% (Fig. 2E) and significantly delayed disease progression (Fig. 2F).

### Targeting TOPORS functionally spares healthy hematopoiesis

To assess the combination’s relative toxicity to healthy cells, we used a hybrid lentiviral/electroporation CRISPR gene-editing strategy^34^ to edit *TOPORS* in human cord blood CD34^+^ HSPCs (Fig. 3A), and confirmed effective polyclonal editing of the *TOPORS* locus by indel frequency analysis (Fig S3A). We then treated sgTOPORS-1 edited CD34^+^ HSPCs with AZA or vehicle for five consecutive days in co-culture with murine MS5 stromal cells and carried out colony forming assays to assess HSPC function. *TOPORS*-editing alone or in combination with AZA therapy did not impair colony forming capacities compared to controls (Fig. 3B).

**Figure 3.**
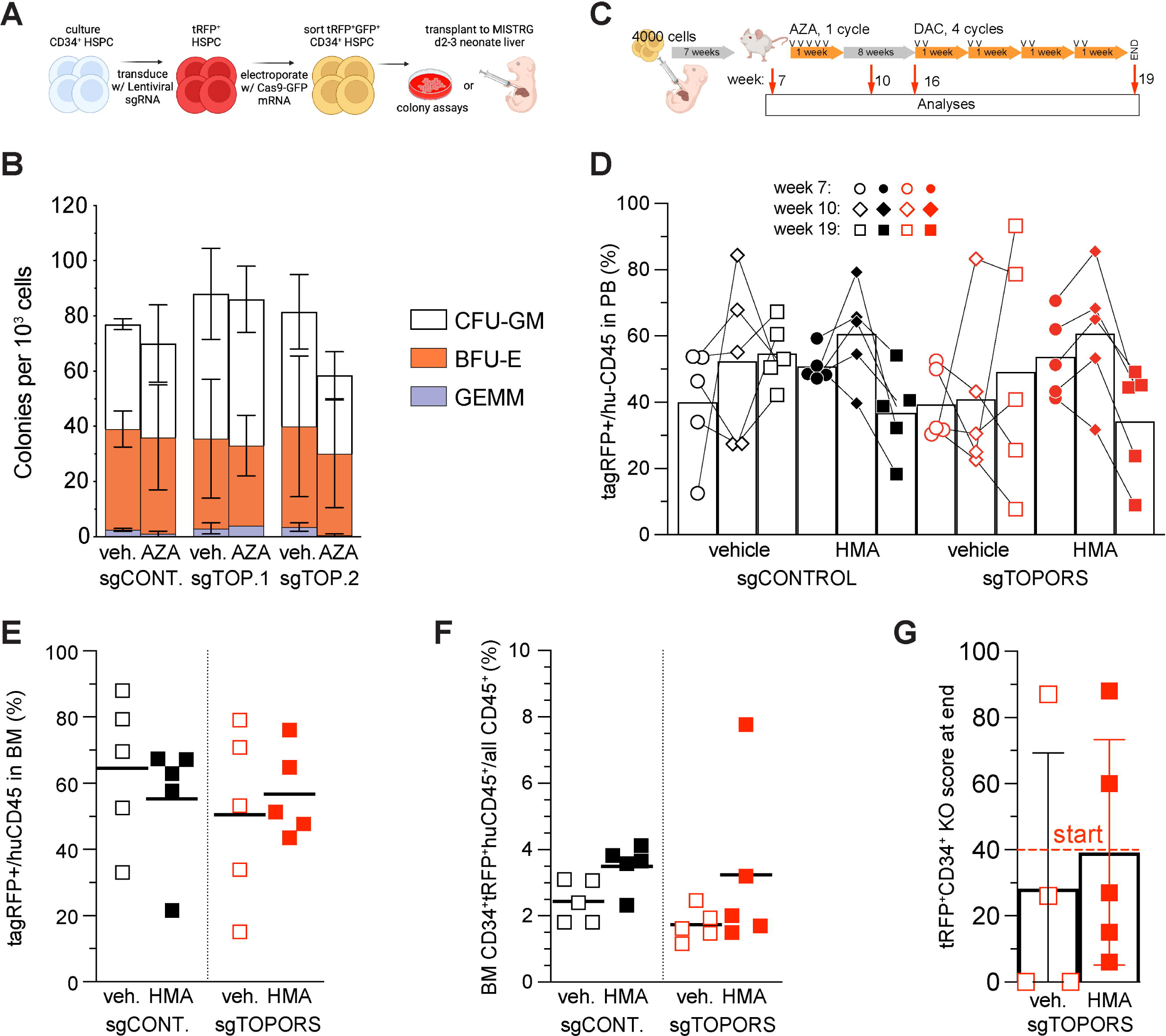
Targeting TOPORS functionally spares healthy hematopoiesis. (A) Workflow for lentiviral stable sgRNA + tagRFP delivery, followed by electroporation of transient Cas9-GFP fusion mRNA into primary CD34^+^ cord blood-derived HSPCs for target gene editing. Image made with BioRender. (B) Colony forming capacity of tagRFP-sorted gene-edited CD34^+^ HSPCs pre-treated with AZA. Data are mean and range; n = 2 experiments using independent cord blood donors. (C) Workflow for engraftment into MISTRG neonates with tagRFP-sorted gene-edited CD34^+^ HSPCs, followed by drug-treatment, blood monitoring, and endpoint analysis of the engrafted adults. Image made with BioRender. (D) Tracking of tagRFP^+^-frequencies in huCD45^+^moCD45^-^ cells for each engrafted mouse indicating frequencies (circles) before drug treatment, (diamonds) after one AZA cycle, and (squares) after AZA followed by DAC cycles. Bars indicate means for each treatment group (n = 5 per group). (E-F) Endpoint bone marrow frequencies for each engrafted mouse, with line for mean; (E) % tagRFP^+^ cells amongst huCD45^+^moCD45^-^ cells, or (F) % CD34^+^ tagRFP^+^ huCD45^+^ moCD45^-^ cells amongst all CD45^+^ cells. (G) Polyclonal indel KO scores generated by ICE algorithm for tagRFP^+^CD34^+^huCD45^+^moCD45^-^ cells sorted from endpoint bone marrows. Bars: means ± SD. Dashed line: Polyclonal indel KO score generated by ICE algorithm for day of engraftment.

To assess the toxicity of targeting TOPORS *in vivo*, we transplanted *TOPORS*-edited cord blood CD34+ HSPCs into neonatal MISTRG recipients^33^ (Fig. 3C) and monitored engraftment by tracking peripheral blood once the transplanted mice had reached adulthood, and again after administering AZA at 1mg/kg for five consecutive days (a single treatment cycle). Neither *TOPORS*-editing alone nor in combination with AZA treatment significantly reduced the frequencies of circulating cells, relative to the non-targeting control (Fig. 3D), although we did note long-term selection against engineered cells (i.e. tagRFP^+^ cells) in favour of non-engineered (i.e. tagRFP^-^) human cells irrespective of the sgRNA (Fig. S3B). After 8 weeks of recovery, we evaluated the combination of *TOPORS*-editing and DAC-treatment (co-administered with the cytidine deaminase inhibitor tetrahydrouridine [THU]), on the same engrafted mice, by treating with 10mg/kg THU i.p. plus 0.1mg/kg DAC or vehicle s.c. for two consecutive days per weekly cycle for a total of four cycles. This low dose and frequency regimen, which minimizes metabolic adaptation and prolongs HMA-sensitivity in the clinic^3,5^, did not affect the production of *TOPORS*-edited blood cells relative to control cells (Fig. 3D, Fig. S3B), nor did *TOPORS*-editing impact the survival of human CD34+ progenitor cells in mouse bone marrow with or without sequential drug treatments (Fig. 3E-F). Although variable between mice, the mean indel polyclonal KO scores for *TOPORS*-edited human CD34^+^ cells exposed to HMA *in vivo* remained comparable to the pre-treatment score, suggesting that *TOPORS* editing does not confer a selective disadvantage on healthy human CD34^+^ cells (Fig. 3G). Unexpectedly, *TOPORS*-editing may have biased towards accumulation of CD56+ NK cells in bone marrow of endpoint mice regardless of HMA therapy, although this trend was not significant (Fig. S3C).

### Targeting TOPORS primes HMA response in leukemic cells via deficient DDR

TOPORS, an Arg/Ser-(RS-) rich ring finger domain protein bearing SUMO interaction motifs (SIM), was the first discovered dual ubiquitin and SUMO E3-ligase^28,30^. TOPORS activates DNA repair pathways by SUMOylating DNA repair factors and chromatin proteins, including TP53, and down regulates the same pathways and overlapping factors *via* ubiquitination; TOPORS is itself a SUMOylation substrate^35–37^.

We evaluated whether TOPORS participates in HMA-induced DDR by measuring ψH2AX levels, a marker of DNA breaks, in AZA-treated MDS-L cells. Compared to control cells, *TOPORS*-edited cells accumulated significantly more γH2AX in response to AZA (Fig. 4A), indicating more DNA break persistence. This conclusion was supported by “comet” assays, where DNA fragments from AZA-treated *TOPORS*-edited MDS-L cells migrated during electrophoresis with larger tail moments compared to control cells (Fig. 4B). Flow cytometry of fixed cells revealed that AZA-treated *TOPORS*-edited MDS-L cells accumulated in late S- and/or G2/M phases (Fig. 4C–D), suggesting that AZA-induced DNA damage delayed cell cycle progression of TOPORS-deficient cells subsequent to incorporation of 5 aza-dC into DNA. Parallel annexin V co-staining revealed that AZA treatment triggered higher levels of apoptosis in *TOPORS*-edited MDS-L cells compared to control cells (Fig. 4E).

**Figure 4.**
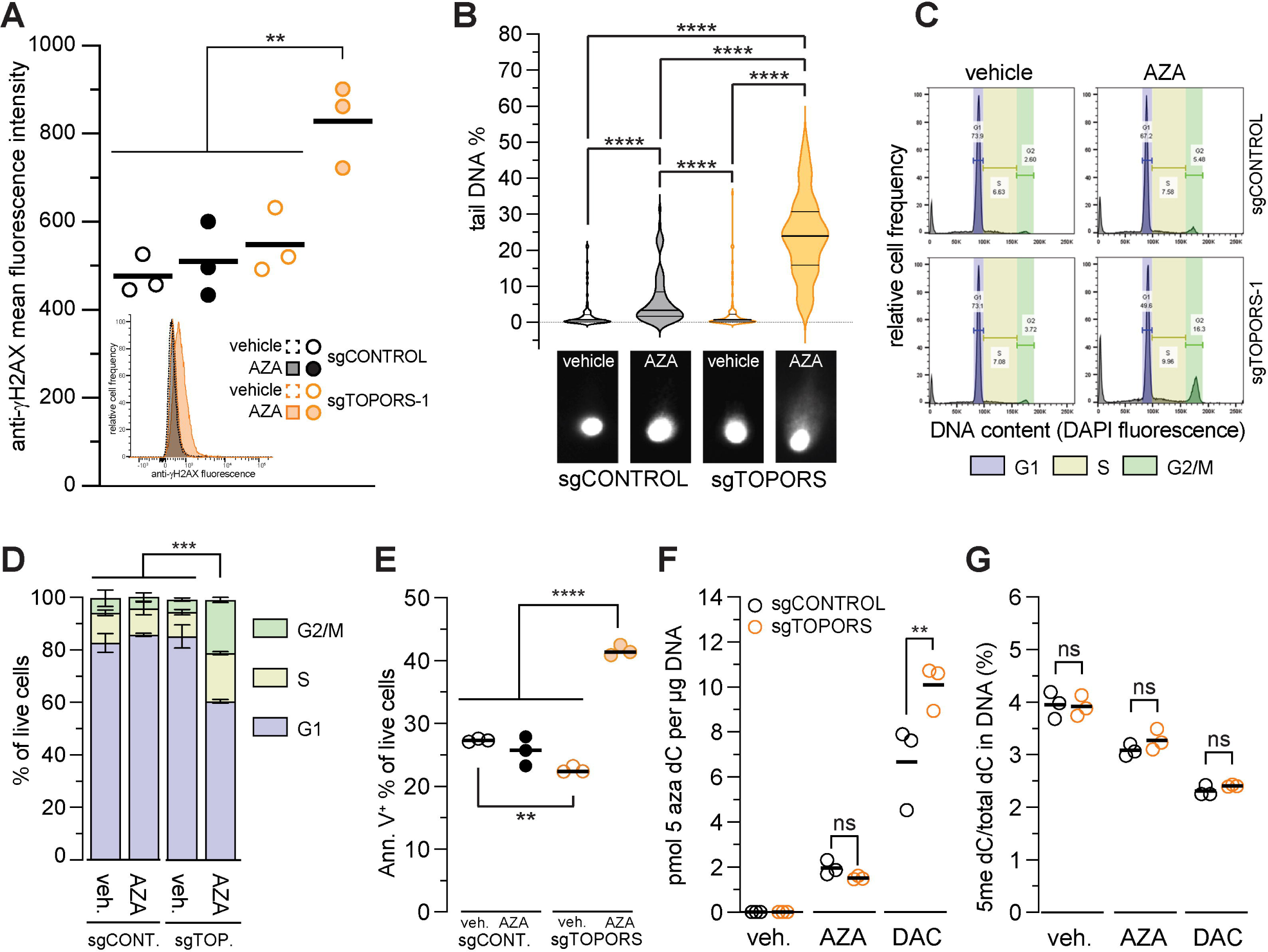
Targeting TOPORS sensitizes leukemic cells to HMAs *via* defective DDR. (A) Mean fluorescence intensity of anti-γH2AX staining by FACS of fixed/permeabilized gene-edited MDS-L cells treated daily with 0.3μM AZA for 4 days. ** P < 0.01 compared to every other treatment by one-way ANOVA, n = 3; P > 0.05 comparisons not shown. Inset: FACS data representing the middle data point for each treatment. (B) Detection of DNA breaks by comet assay in gene-edited MDS-L cells treated with 0.3μM AZA as for A. **** P ≤ 0.0001, for Kruskal-Wallis multiple comparison tests, n = 75; P > 0.05 are not shown. (C) Example DNA content profiles determined by DAPI staining of gene-edited MDS-L cells treated with 0.3μM AZA. (D) Stacked histograms from biological triplicates of C; *** P < 0.001 compared to every other treatment by two-way ANOVA; P > 0.05 comparisons not shown. (E) Proportion of apoptotic cells determined by Annexin/PI staining in gene-edited MDS-L cells treated with 0.3μM AZA. **** P < 0.0001, ** P < 0.01, one-way ANOVA, n = 3; P > 0.05 comparisons not shown. (F) Incorporation of 5 aza-dC, and (G) methylation at dC, in genomic DNA (both determined by LC-MS) in gene-edited MDS-L cells exposed to 0.3µM AZA or 0.02µM DAC daily as for A. ** P ≤ 0.01, ^ns^ P > 0.05, for selected pairwise comparisons in one-way ANOVA, n = 3.

Because HMA incorporation into DNA is S-phase dependent, we used mass spectrometry of digested DNA to test whether cell cycle changes induced by editing *TOPORS* might alter either incorporation of 5 aza-dC into DNA, or subsequent DNA demethylation^38^. *TOPORS*-editing did not significantly change incorporation of 5 aza-dC into DNA of cells treated with AZA but led to increased incorporation into DNA of cells treated with DAC (Fig. 4F). Genome demethylation in response to 5 aza-dC incorporation was similar in *TOPORS*-edited versus control cells (Fig. 4G).

### Multi-omic approaches reveal widespread mis-splicing of DDR genes and cycle arrest in AZA-treated TOPORS-edited MDS-L cells

Besides its role in post-translational modification of proteins, TOPORS has also been characterized as a transcriptional regulator through binding *cis-* regulatory elements and influencing chromatin accessibility at enhancers^39,40^. As these findings suggest that TOPORS and AZA potentially converge through epigenetic remodelling and subsequent changes in gene expression, we assessed the bulk transcriptomes and nuclear proteomes of *TOPORS*-edited MDS-L cells treated with AZA. Gene set enrichment analyses revealed marked enrichment of DNA replication and spliceosome transcriptional programs in AZA-treated *TOPORS*-edited MDS-L cells compared to controls (Fig. 5A). Inference of transcriptional regulatory networks underlying these alterations using the TRRUST database^41^ identified that E2F1 targets significantly overlap upregulated genes in AZA-treated *TOPORS*-edited MDS-L cells (Fig. 5B). Clustering of transcriptomes based on expression of E2F1 targets revealed that AZA-treated *TOPORS*-edited MDS-L transcriptomes displayed the greatest enrichment of E2F target genes (Fig. S4A). E2F1 is a transcription factor induced in response to DNA damage, with major roles in cell cycle progression and DNA repair^42^. E2F1 localizes to sites of DNA damage and origins of replication to collaborate with DNA repair proteins, enabling DNA repair and completion of DNA replication^43^. Thus, increased DNA replication programs in AZA-treated *TOPORS*-edited MDS-L cells appear to be driven by increased E2F1 activity.

**Figure 5.**
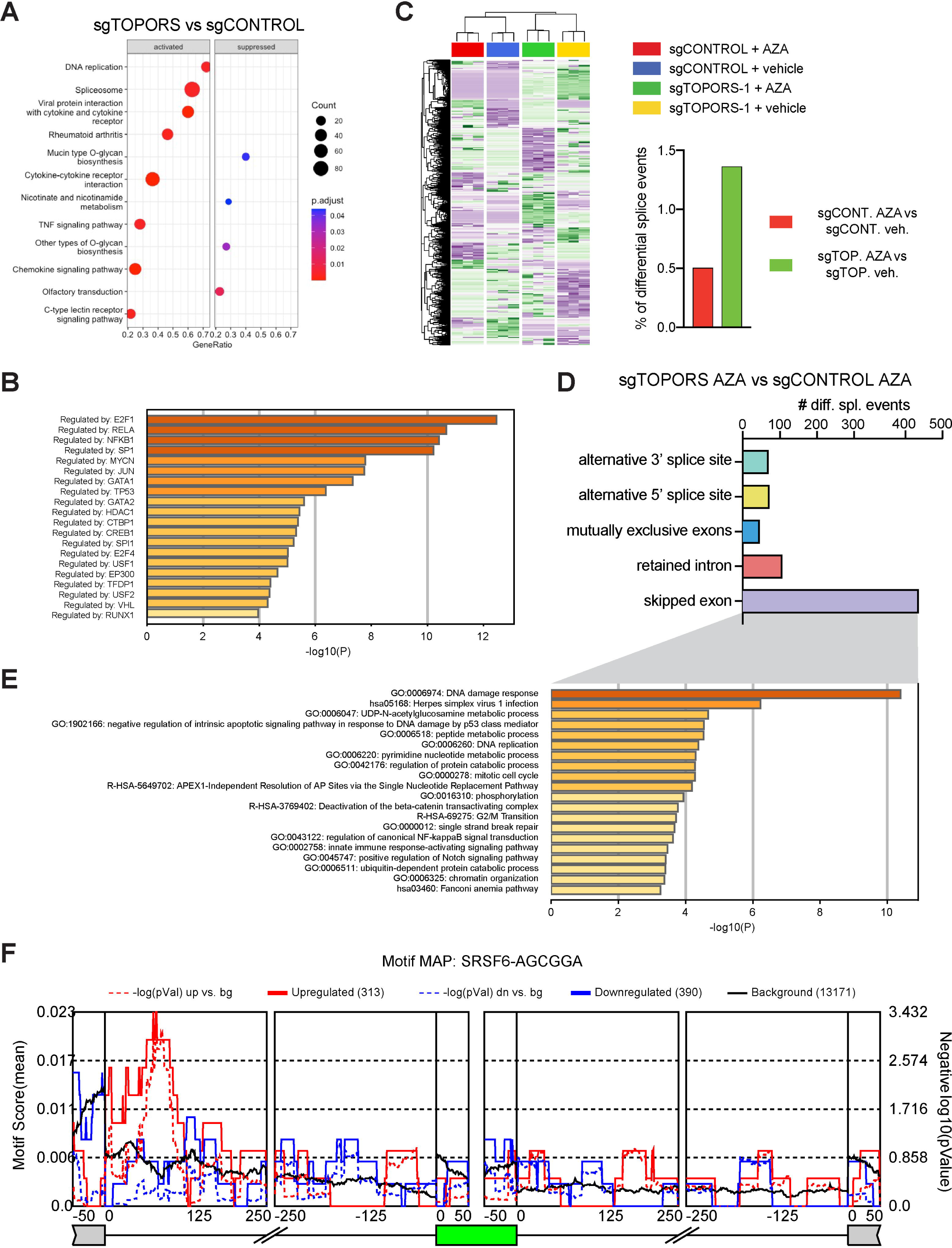
Multi-omic approaches reveal widespread mis-splicing of DDR genes and cycle delay in AZA-treated *TOPORS*-edited MDS-L cells. (A) Gene set enrichment analysis of differentially expressed genes using the KEGG module as part of the clusterProfiler algorithm. Dot plots depict gene sets that are enriched or suppressed in AZA-treated *TOPORS*-edited MDS-L cells compared to AZA-treated control cells. GeneRatio refers to the ratio of input genes that are annotated in a term. (B) Transcription factors with targets demonstrating significant overlap with differentially upregulated genes in AZA-treated *TOPORS*-edited MDS-L cells compared to vehicle determined through TRRUST. (C) Unsupervised hierarchical clustering and heatmap of 3826 most significant differentially spliced events between all samples (n=12). Histograms at right show the proportion of mis-spliced events detected as percentage of all splicing events. (D) Number of alterative splicing events detected in AZA-treated *TOPORS*-edited MDS-L compared to AZA-treated control cells. (E) Overrepresentation analysis of skipped exon events detected in AZA-treated *TOPORS*-edited MDS-L cells compared to AZA-treated control cells. (F) Motif scanning analysis for AGCGGA (SRSF6) binding sites across a meta-exon (green) generated from all exon skipping events. Motif enrichment scores (left axis) and -log_10_(P values) (right axis) are shown. (red) Motif enrichment scores of exons differentially retained in AZA-treated *TOPORS*-edited MDS-L cells. (blue) Motif enrichment scores of exons differentially skipped in AZA-treated *TOPORS*-edited MDS-L cells. (dashed) Significance scores. (black) background score calculated from all non-differentially spliced exons.

Chemotherapeutics that induce DNA-adducts impair transcription through steric hindrance^44^. Bulky lesions in particular cause transcription-dependent splicing alterations^45^ that result in use of weaker splice sites and facilitates the inclusion of suboptimal exons^46^. Therefore, AZA induced formation of bulky DNA-DNMT1 adducts could potentially impact alternative splicing. Alternative splicing alterations can also be triggered by variations in expression levels or post-translational modifications of splicing factors^47^. According to the BioPlex human interactome database^48^, TOPORS interacts with splicing factors SRSF4 and SRSF6 (Fig. S4B) and has been shown to SUMOylate other splicing factors^31^. For these reasons, along with the enrichment of spliceosome signatures in AZA-treated *TOPORS*-edited MDS-L cells, we investigated the alternative splicing landscape of these cells. We used rMATs^49^ to quantify five alternative splicing events from our bulk transcriptomic datasets. Unsupervised hierarchical clustering and principal component analysis of all samples revealed that AZA treatment and *TOPORS*-editing resulted in global splicing alternations both individually and cooperatively (Fig. 5C).

We detected 764 mis-spliced events in AZA-treated *TOPORS*-edited MDS-L cells compared to control cells, the majority being exon skipping events (n=473, FDR <0.05, PSI>0.1) (Fig. 5D). Over-representation analysis associated these mis-spliced genes with pathways relating to the DDR (Fig. 5E). To test whether specific RNA binding proteins were responsible for these exon skipping events, we used rMAPs^50^ to assess the density of RNA binding protein motifs within the associated sequences. We identified significant enrichment for a motif (AGCGGA) bound by SRSF6, a spliceosome factor known to interact with TOPORS in HEK293 cells^48^. Exons 3’ to this motif were differentially retained in AZA-treated *TOPORS*-edited MDS-L cells (Fig. 5F). We also identified significant enrichment for two SRSF1 binding motifs but did not identify any specific enrichment for SRSF4 motifs (Fig. S4C). Defective SRSF6-dependent DDR transcript splicing might therefore be a contributing factor to the AZA sensitivity of *TOPORS*-edited MDS-L cells.

Since TOPORS is primarily nuclear-localized, we profiled the nuclear proteome of AZA-treated and *TOPORS*-edited MDS-L cells using label-free mass spectrometry. On average, 1500 proteins were detected per sample with a minimum coverage of 1000 proteins between samples (Fig. S5A). Similar to our transcriptomics data, AZA-treated *TOPORS*-edited cells had a distinct nuclear proteome compared to controls (Fig. S5B). Differential abundance analysis identified 73 of 2257 proteins as significantly differentially abundant across all samples (Fig. 6A). After hierarchical clustering, cluster 1 represented proteins that were down-regulated specifically in steady state cells by TOPORS activity – presumably via TOPORS-mediated ubiquitination. Clusters 4 and 6 represented proteins that were enriched or depleted, respectively, in AZA-treated *TOPORS*-edited MDS-L cells (Fig. 6A). Over-representation analyses indicated that cluster 4 proteins were associated with late-stage cell cycle proteins, while cluster 6 proteins were associated with global nucleotide excision repair pathways (Fig. 6B). DNMT1 (bottom of cluster 5) was significantly depleted in AZA-treated MDS-L cells, regardless of *TOPORS* editing (Fig. 6B). This was also so in DAC-treated MOLM-13 cells (Fig. 6C), indicating that TOPORS-deficiency did not prevent HMA-induced depletion of DNMT1. The notably higher levels of protein SUMOylation we observed in DAC-treated *TOPORS*-edited MOLM13 cells (Fig. 6C), indicated that TOPORS was not needed for DNA damage-induced SUMOylation generally. Nonetheless, our finding that DNMT1 levels in AZA-treated *TOPORS* edited cells were two-fold higher compared to the AZA control (t-test p=0.03; Fig. 6B), indicated that AZA-induced DNMT1 degradation might be specifically influenced by TOPORS activity. Overall, our transcriptomic and proteomic findings were consistent with the impaired DNA damage and cell cycle arrest signatures identified in functional assays and indicated that TOPORS-deficiency might synergize with AZA both by impeding the removal of AZA-induced DNMT1-DNA adducts and by altering RNA splicing during AZA-induced DDR.

**Figure 6.**
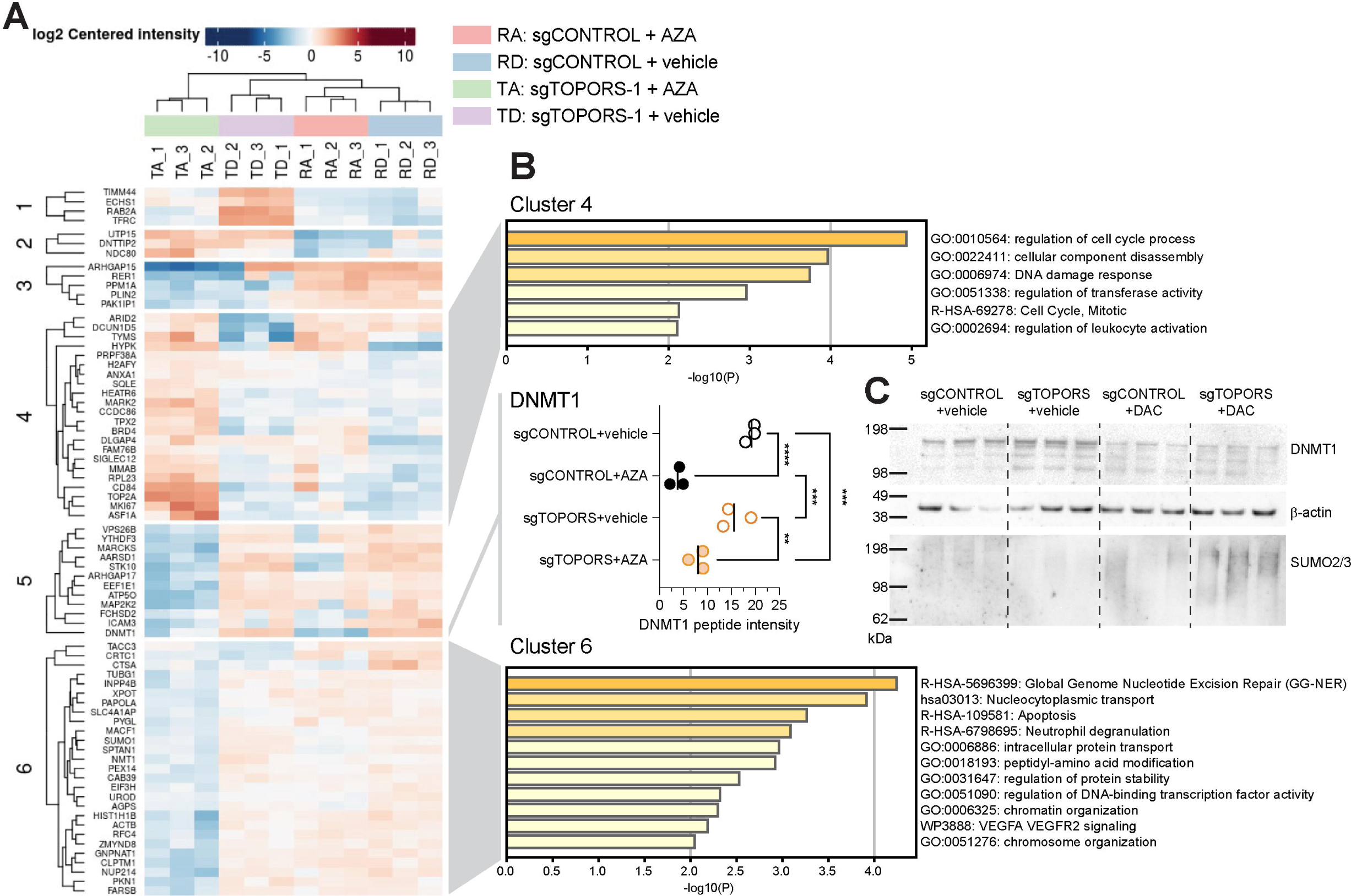
Nuclear proteomics reveals a depletion of global nucleotide excision repair factors in AZA-treated *TOPORS*-edited MDS-L cells. (A) Heatmap of unsupervised hierarchical clustering of 73 proteins that were significantly differentially abundant across all nuclear extracts from triplicate cultures of gene-edited MDS-L cells treated daily with 0.3µM AZA or vehicle for 4 days; n = 12. (B) Overrepresentation pathway analysis of proteins enriched or depleted in (top) cluster 4 and (bottom) cluster 6. (middle) Normalized total spectra (Scaffold 5.3.2) for DNMT1 peptides detected in each replicate. **** P < 0.0001, *** P < 0.001, ** P < 0.01, one-way ANOVA, n = 3; P > 0.05 comparisons not shown. (C) Sequential detection in the same western blot of (top) DNMT1, then (middle) β-actin, then (bottom) SUMO2/3 in 10µg nuclear proteins from gene-edited MOLM-13 cells treated with 42nM DAC or vehicle daily in triplicates for 3 days.

### TOPORS-editing sensitizes cells to HMA in a DNMT1-dependent manner

To formally test whether genome demethylation *per se* or DNMT1 were involved in HMA-hypersensitivity in *TOPORS*-edited cells, we measured sensitivity to a novel DNMT1 small molecule inhibitor (GSK3685032) which inhibits DNMT1 without incorporating into nucleic acids ^51^. *TOPORS*-edited cells were not more sensitive to GSK3685032 than control cells, even at GSK3685032 concentrations inducing very high levels of cytidine demethylation (Fig. 7A). We formally tested whether DNMT1 was involved in MDS-L HMA-sensitivity by exposing cells to DAC in the presence or absence of 1µM GSK3685032, a concentration which induced near-maximal DNMT1 inhibition with minimal cytotoxicity (Fig. 7A, Fig. 7B). DAC-mediated killing of both *TOPORS*-edited and control MDS-L cells was reduced equally (by 7.4-fold; Table S3) in the presence of 1µM GSK3685032, implicating DNMT1 adduction as the dominant cytotoxic event in DAC-treated MDS-L cells, regardless of TOPORS activity. To test whether other replication-blocking DNA-protein adducts require TOPORS for efficient resolution, we assessed sensitivity to the topoisomerase inhibitors topotecan or etoposide, which prevent type 1 or type 2 topoisomerases from resolving the transient tyrosine-DNA ester bonds they form with DNA. *TOPORS*-editing did not sensitize MDS-L cells to either topoisomerase I or II inhibition by a single dose of topotecan or etoposide (Fig. 7C-D), and furthermore did not sensitize to hydroxyurea – a ribonucleotide reductase poison that disrupts nucleotide synthesis (Fig. 7E). Thus, TOPORS-deficiency sensitizes to DNA-DNMT1 adducts specifically, and not to disrupted nucleotide metabolism nor to DNA-protein adducts in general.

**Figure 7.**
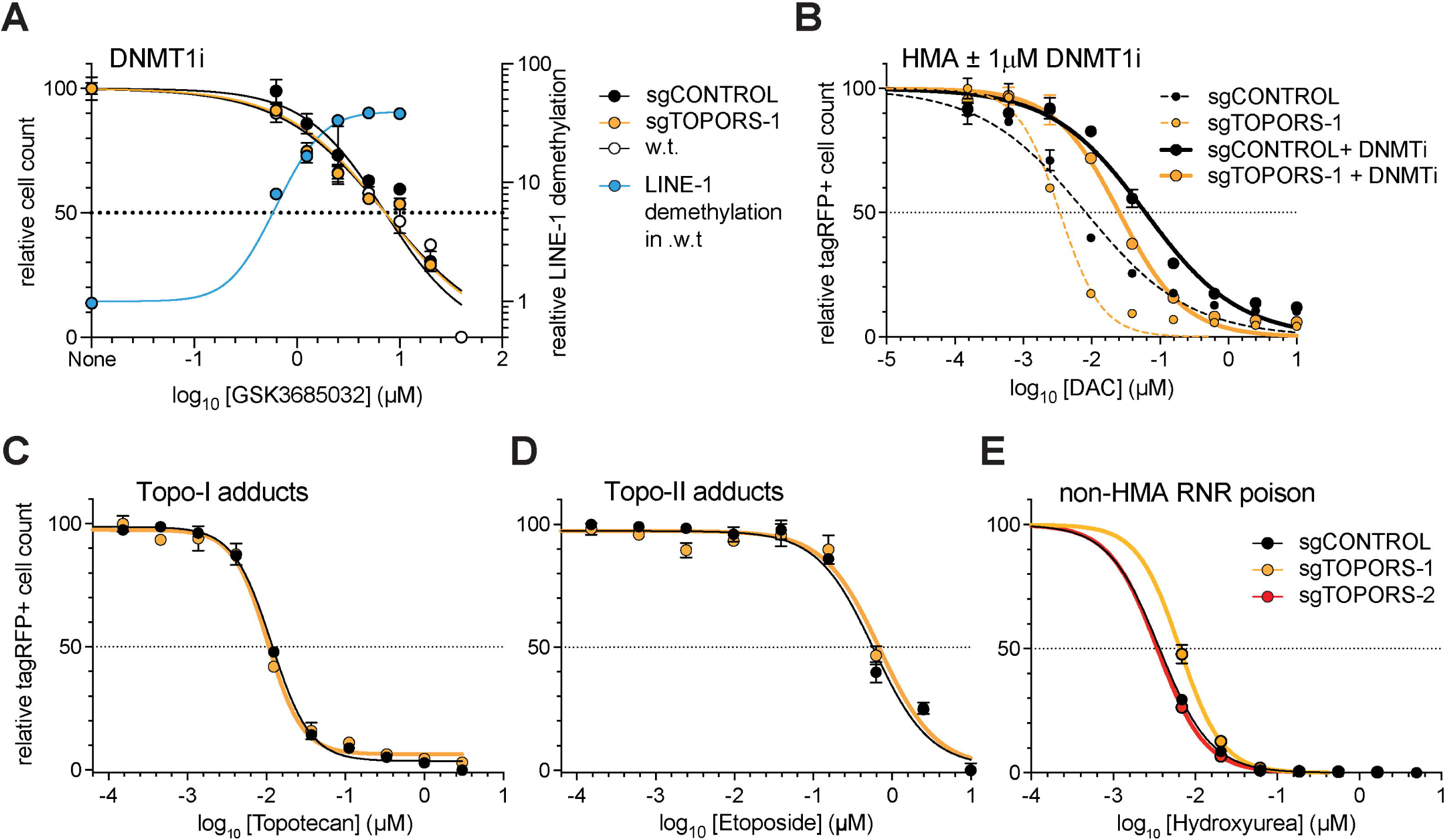
*TOPORS-*editing sensitizes cells to HMA in a DNMT1-dependent manner. (A) (left axis) Survival of gene-edited or wild-type MDS-L cells in 96-well plates in response to daily doses of DNMTi GSK3685032 or vehicle on days 1–4. (left axis) tagRFP+ cells (or all cells for wild-type MDS-L) were counted on day 5 and normalized to vehicle counts. (right axis) In a separate experiment, relative demethylation of LINE-1 promoters in wild-type MDS-L was determined (see Methods) for a similar GSK3685032 dose range and normalized to vehicle control cells. (B) Gene-edited MDS-L cells were plated into 96-well plates and treated with 1µM GSK3685032 or vehicle on day 1. Varying DAC plus 1µM GSK3685032 or vehicle was added daily on days 2–4, and tagRFP+ cells counted on day 5, and normalized to vehicle counts. (C-D) Gene-edited MDS-L cells were plated into 96-well plates and treated day 1 only with (C) Topotecan or (D) Etoposide or vehicle. tagRFP+ cells were counted on day 5 and normalized to vehicle counts. (E) Gene-edited MDS-L cells in 96-well plates were treated with daily Hydroxyurea or vehicle on days 1–4. tagRFP+ cells were counted on the day 5 and normalized to vehicle counts. In all panels, dots represent mean ± SD, (A–B) n = 3, (C–E) n = 4. EC50 values deduced from all panels are shown in Table S3.

### TOPORS-editing does not reduce SUMOylation of DNMT1 in HMA-treated AML cells

DNMT1 SUMOylation followed by RNF4-mediated SUMO-targeted ubiquitylation are critical for clearance of HMA-induced DNMT1 adducts^52,53^. This lead us to suspect that TOPORS may mediate DNMT1 SUMOylation, but the potential importance of interactions between TOPORS and other targets, such as RNA-splicing factors, prompted us to explore TOPORS’ E3 SUMO-ligase activity in an unbiased manner in AML cells. *TOPORS*-edited or control MOLM-13/Cas-9 cells were engineered to co-express Dasher-GFP plus SUMO1 N-terminally tagged with 10xHis (10His-SUMO1, Fig. S6A). Sorted Dasher-GFP+ cells were treated for 3 consecutive days with 4.2 nM DAC or vehicle then whole cell guanidine-extracted proteins bearing 10xHis tags were enriched by pull-down with Ni-NTA beads^54^, and subjected to label-free LC-MS/MS analyses. Mass spectral yields for 10xHis-enriched proteins were substantially lower and more variable between replicates than for our prior nuclear proteomics (Fig. S5; Fig. S6B–C). Nonetheless, a small set of Ni-captured proteins, including DNMT1, were significantly differentially abundant across the four experiment conditions, (Fig. S6D). One of the highest ranked Ni-captured proteins depleted from *TOPORS*-edited cells, regardless of DAC treatment, was TOPORS itself (Fig. S6D, Fig. 8A– C). This unequivocally confirmed that *TOPORS*-editing had substantially depleted TOPORS protein and/or its E3 SUMO-ligase activity from AML cells. Other proteins differentially captured by Ni-NTA from vehicle-treated (steady state) cells were likely E3-ligase targets of TOPORS; these were predominantly involved in ribonucleoprotein complex biogenesis and in RNA splicing (Fig. 8D), which was strikingly consistent with our prior transcriptome and nuclear proteome datasets. Focusing on DNMT1, SUMOylated DNMT1 was almost equally Ni-NTA captured from DAC-treated control and TOPORS-edited cells, but not from vehicle-treated cells (Fig. 8C). Since total DNMT1 levels drop substantially in MDS/AML cells chronically treated with low dose HMA, regardless of TOPORS activity (Fig. 6B-C), we deduce that the SUMOylation level of the remaining DNMT1 was very high, and that TOPORS is not required for HMA-mediated DNMT1 SUMOylation. Indeed, our prior orthogonal experiments indicated that the levels of SUMOylated DNMT1 were higher in HMA-treated TOPORS-edited compared to controls. Furthermore, our combined data indicate that reduced TOPORS-mediated SUMOylation or ubiquitylation of proteins involved in RNA metabolism and splicing (Fig. 8D) likely plays an important role in the sensitivity of *TOPORS*-edited cells to HMA.

**Figure 8.**
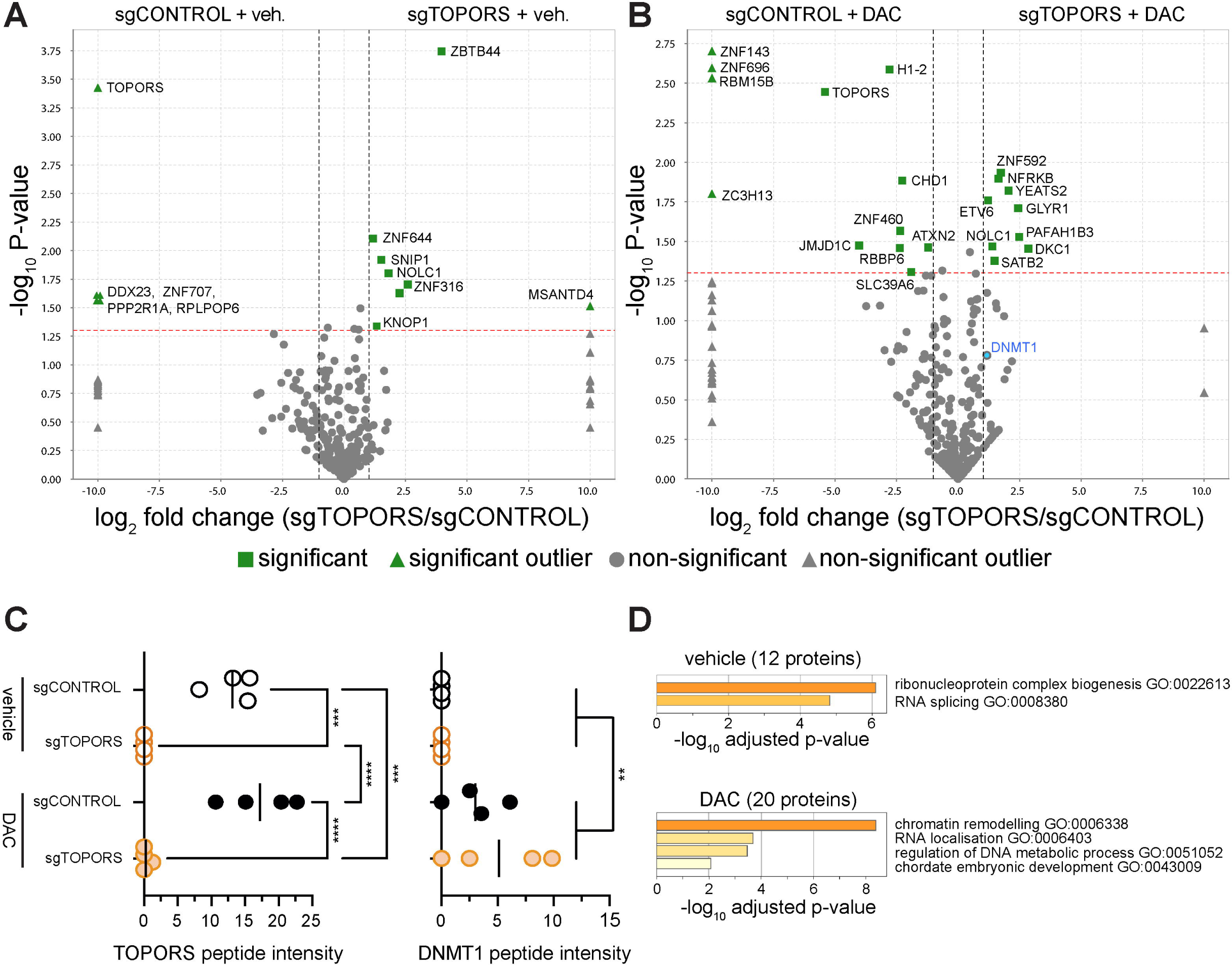
*TOPORS-*editing does not reduce SUMOylation of DNMT1 in HMA-treated AML cells. Proteomic analysis of whole cell Ni-NTA enriched proteins from gene-edited MOLM-13 cells expressing 10xHis-SUMO1 that were exposed to 42 nM DAC or vehicle for 3 days. (A–B) Volcano plots (Scaffold 5.3.2) highlighting proteins significantly more that 2-fold enriched in *TOPORS*-edited cells compared to control cells under (A) “steady-state” (i.e. vehicle treated) conditions or (B) after exposure to DAC. (C) Normalized total spectra (Scaffold 5.3.2) for (left) TOPORS or (right) DNMT1 peptides in each replicate. **** P < 0.0001, *** P < 0.001, ** P < 0.01, by (TOPORS) one-way ANOVA or (DNMT1) t-test; P > 0.05 comparisons not shown. (D) Summary of enrichment into GO Biological pathways for Ni-NTA captured proteins that were differentially abundant between *TOPORS*-edited versus control cells for (top) vehicle-treated or (bottom) DAC-treated conditions.

### SUMOylation blockade synergizes with HMAs in MDS and AML in vivo

Concurrent with pursuing the mechanistic basis of synergy between *TOPORS*-editing and HMAs, we evaluated whether our findings are translatable to clinical application. There are currently no pharmacological molecules specifically inhibiting TOPORS. However, the prior known importance of SUMOylation in clearing HMA-induced DNMT1 adducts suggested that inhibiting SUMOylation might enhance the potency of HMA therapy. The first-in-class drug TAK-981 acts upstream of SUMO E3-ligases such as TOPORS by preventing attachment of SUMO to the universal SUMO E2-conjugase UBC9, which leads to global inhibition of protein SUMOylation^55^. Combinatorial treatment with TAK-981 and AZA was cytotoxically additive in MDS-L and Kasumi-1 (both *TP53* mutant), but synergistic in MOLM-13 and TF-1 cell lines (both *TP53* wild-type, Fig. 9A and Fig. S7A). Critically, TAK-981 and DAC combination was synergistic in all four cell lines (Fig. 9B and Fig. S7B). Lower synergism with AZA may reflect SUMO-independent activities mediated by drug incorporation into RNA, which are not replicated by DAC. We also measured the impact of low concentration TAK-981 (0.1µM) on HMA cytotoxicity in *TOPORS*-edited versus control MDS-L cells. The difference in HMA-sensitivity between control and *TOPORS*-edited cells was much smaller in the presence of TAK-981 (AZA:1.2-fold, DAC: 1.4-fold) compared to its absence (AZA: 1.99-fold, DAC: 1.94-fold Fig. 9B, Fig. 9C, Table S4), demonstrating that TOPORS mediates a substantial proportion of SUMO-dependent survival in HMA-exposed MDS-L cells. To determine whether TAK-981 could phenocopy the DDR-deficient signature induced by *TOPORS*-editing, we treated MDS-L with the lowest synergistic dosage of TAK-981 and HMAs (from synergy maps in Fig. S7) and assessed γH2AX levels and cell cycle status. Low-dose TAK-981 combined with low-dose HMA resulted in extensive accumulation of γH2AX with concomitant accumulation of cells in the late S and G2/M phases (Fig. 9D-E).

**Figure 9.**
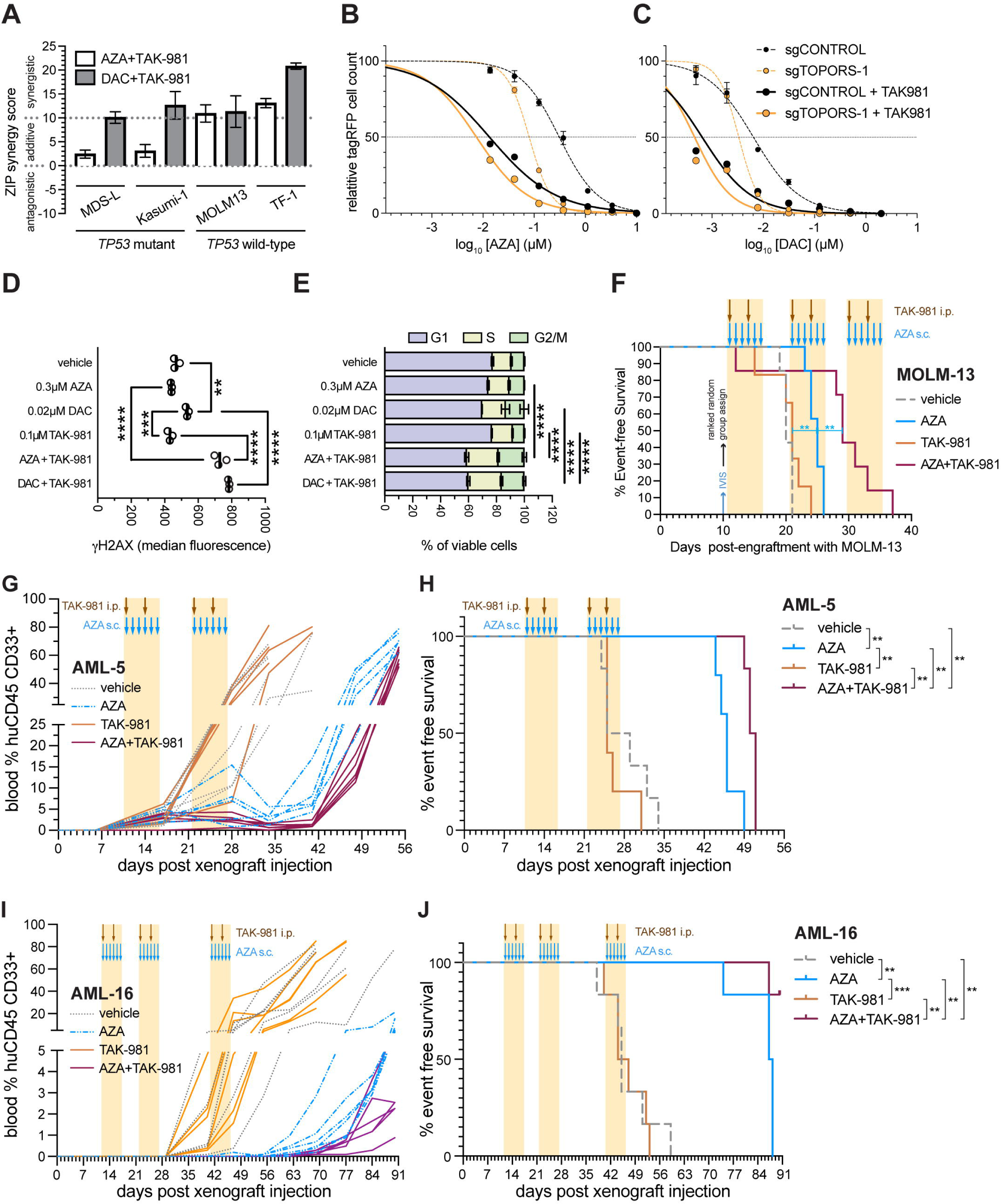
SUMOylation blockade synergizes with HMAs in MDS and AML. (A) Summary of ZIP synergy scores (± 95% CI) for combinatorial drug testing in MDS-L and AML lines determined by SynergyFinder. (B-C) Survival of gene-edited MDS-L cells in 96-well plates in response to (B) AZA ±0.1µM TAK-981or (C) DAC ±0.1µM TAK-981 added daily on days 1–4. TagRFP+ cells were counted on day 5, and normalized to vehicle counts ±SD. EC50 values are listed in Table S4. (D) Mean fluorescence intensity of anti-ψH2AX staining by FACS of fixed/permeabilized gene-edited MDS-L cells drug-treated as in B–C. **** P < 0.0001, *** P < 0.001, ** P < 0.01, by one-way ANOVA; P > 0.05 comparisons not shown. (E) Cell cycle distributions ±SD for the same cultures as D (n=3); **** P < 0.0001 by two-way ANOVA; only comparisons between combination and single drugs shown. (F) Event-free survival of non-irradiated MISTRG engrafted i.v. with MOLM-13 cells (n = 5–7 per treatment). Mice were sex and IVIS flux rank randomised in cohorts of 4 into treatment groups on day 10, then drug treatments (20mg/kg TAK-981 i.p., 0.6mg/kg AZA s.c.) commenced on day 11. ** Gehan-Breslow-Wilcoxon test P < 0.01 for AZA versus any other treatment. (G-J) Sub-lethally irradiated MISTRG mice were injected with 1.25×10^6^ AML-5 PDX cells i.v.; non-irradiated MISTRG mice were injected with 4×10^6^ AML-16 PDX cells i.v. PDX-injected mice were then bled at approximately weekly intervals. The mice were randomised in sex and weight ranked cohorts of 4 into treatment groups on day 10 or 11, then treated with the drugs (20mg/kg TAK-981 i.p., 0.6mg/kg AZA s.c.) starting day 11 or 12. (G,I) Spaghetti plots of blood % huCD45^+^CD33^+^ cells, to track expansion of each (G) AML-5 or (I) AML-16 PDX in xenografted mice. The (G) 25% event threshold or (I) 5% event threshold is indicated. (H and J) Kaplan-Meier plots for event-free survival of the same mice as G and I. Event-free survival was scored as time to reach (H) 25% or (I) 5% engraftment of huCD45^+^CD33^+^ cells in peripheral blood ^55^. *** P < 0.001, ** P < 0.01, by Gehan-Breslow-Wilcoxon test; P > 0.05 comparisons not shown.

We next asked whether *in vitro* synergy between TAK-981 and HMAs extended to a more clinically relevant *in vivo* context. TAK-981 combined with low dose AZA synergistically and significantly prolonged survival in MISTRG mice engrafted with MOLM-13 cells (Fig. 9F). To determine whether TAK-981 also sensitizes primary patient AML cells to AZA *in vivo*, we transplanted patient-derived continuous xenograft (PDX) lines AML-5 and AML-16 ^56^ into MISTRG mice, followed by drug treatment (Fig. 9G–J). Expansion of these PDX was significantly delayed by AZA therapy alone, but not by TAK-981 alone, and expansion was synergistically delayed by combined AZA plus TAK-981 therapy (Fig. 9G–J).

Finally, to test the impact of combined TAK-981 and AZA treatment on healthy cells we injected escalating doses of TAK-981 i.p., combined with AZA s.c., into adult MISTRG mice engrafted with cord blood CD34^+^ cells (Fig. S8A). The frequencies of huCD45^+^moCD45^-^ cells amongst all CD45^+^ cells in MISTRG bone marrow were determined by cytometry at endpoint and revealed no significant differences between treatment groups in endpoint levels of human CD45^+^ cell persistence (Fig. S8B). Thus, combined TAK-981 and HMA treatment is preferentially effective against leukemic cells whilst sparing healthy blood stem and progenitor cells *in vivo*.

## DISCUSSION

In this study, we established a role for the dual E3 ubiquitin and SUMO ligase TOPORS as a central regulator of the DDR induced by incorporation of 5 aza-dC into DNA. We provide genetic proof of concept that targeting TOPORS confers hypersensitivity to HMAs through predisposing leukemia cells to a defective response to DNA-DNMT1 adducts. This hypersensitive phenotype was not dependent on the *del5q* or *TP53* mutational status. Furthermore, we demonstrate that these therapeutic benefits extended to an *in vivo* AML model and that TAK-981 is a viable pharmacological surrogate to targeting *TOPORS*. Our work taken together with reports published during the course of this research suggests that TAK-981 combined with HMAs could offer therapeutic benefits to HR-MDS and AML patients^58^.

By integrating functional and multi-omic approaches, we show that both *TOPORS*-editing and TAK-981 treatment prime an HMA sensitivity phenotype through impairing the DDR to HMA-induced DNA-DNMT1 adducts. Synergy of TAK-981 was higher with DAC compared to AZA, especially in *TP53*-mutant cells. Although TAK-981 is an effective surrogate for TOPORS-editing and demonstrates cytotoxic synergism in combination with low dose HMAs, it should be noted that reduction in TOPORS activity synergizes with HMAs much more specifically than reduction in upstream SUMOylation activity mediated by TAK-981. Complete inactivation of SUMOylation is lethal, while inactivation of E3 ligase TOPORS alone is not^59^. Indeed, in the absence of AZA, MDS-L proliferation in our initial CRISPR-Cas9 screen was reduced in *UBC9*-edited, but not in *TOPORS*-edited cells, while proliferation in the presence of AZA was highly dependent on both UBC9 and TOPORS activities (Table S2). Thus, TOPORS is an attractive target for development of specific inhibitors with potentially improved therapeutic index compared to TAK-981.

Beyond its extensive role in modulating the DDR, SUMOylation plays a crucial role in transcriptional repression of inflammatory cytokines via its influence on chromatin architecture^60^. TAK-981 monotherapy was shown to promote inflammatory anti-tumor immune responses through re-activation of a type I interferon response, and potentiating response to immune checkpoint blockade^55^. These findings align with previous reports where HMA treatment of solid tumors triggered de-repression of silenced endogenous retroviral elements resulting in a potent anti-tumor inflammatory response^17,18^. However, it is unlikely that TAK-981 and HMAs are converging through inflammatory pathways in MDS/AML cells, because these pathways were not enriched in our or other^20,21^ screening datasets, and expression levels of inflammatory pathways do not reliably predict patient response^61^. These findings suggest that either; (1) the level of redundancy in these inflammatory pathways are not conserved between solid tumors and hematopoietic malignancies; (2) the relative expression of these independent inflammatory factors in each model plays a role in HMA dependencies; or (3) these inflammatory pathways primarily modulate HMA response in either a non-proliferative or HSPC-extrinsic manner, readouts that were not captured from our screening efforts.

We confirmed DNMT1 as the dominant mediator of HMA cytotoxicity in AML cells, consistent with previous findings in other tumor models. Against our initial expectations, we found that TOPORS was not required for SUMOylation of adducted DNMT1. However, HMA-induced DNMT1 degradation was nonetheless reduced in *TOPORS*-edited cells. Parallel investigations by others performing similar CRISPR screens identified TOPORS as a SUMO-targeted ubiquitin E3 ligase that acts in semi-redundant concert with RNF4 to mediate efficient proteasomal degradation of DNA-adducted DNMT1^62,63^. Nonetheless, our data strongly indicate that ubiquitylation of SUMOylated DNMT1 adducts is not the only mechanism by which TOPORS protects HMA-exposed cells from DDR-induced apoptosis. Our proteomics identified RNA splicing factors as key candidates for TOPORS-mediated SUMOylation or ubiquitylation in AML cells, even in the absence of HMA, and characterized widespread mis-splicing of DNA repair genes in TOPORS-edited cells, likely due in part to aberrant SUMO- or ubiquitin-modulation of interacting splicing factors. These results draw similarities to a recent study which showed that splicing modulators targeting SF3B1 triggered enhanced exon-skipping in DNA damage repair genes^64^. Widespread mis-splicing of DNA repair genes impaired the DDR in cohesin-mutant AML cells by altering repair protein function, providing a novel approach to sensitize cancer cells to chemotherapeutics and PARP inhibitors. We speculate that targeting TOPORS produces similar deficits in DDR proteins via mis-splicing, and it would be worthwhile to determine whether HMAs could also be combined with splicing modulators for therapeutic benefit.

TOPORS is a promising drug development candidate for HMA combinatorial therapy because TOPORS-editing did not impair S-phase-dependent HMA incorporation into DNA and did not significantly impair blood-forming capacities *in vitro* or *in vivo*. We speculate this was due to greater redundancy between RNF4- and TOPORS-dependent DNA-DNMT1 adduct clearance^53^ in healthy compared to leukemic cells. Our findings that DNMT1 inhibition caused a larger fold-change in DAC-sensitivity in MDS-L cells than inactivation of TOPORS, and that *TOPORS*-editing did not prevent DNMT1-depletion in chronically HMA-treated MDSL/AML cells, demonstrate that such redundancy for DNA-DNMT1 adduct resolution is evident even in leukemic cells.

Our data contrast previous clinical studies where other inhibitors of post-translational mechanisms, including the proteasomal inhibitor Bortezomib (NCT00624936, NCT01420926)^65^ or Pevonedistat (inhibitor of NEDD8 Activating Enzyme) in combination with AZA^66^ did not provide a survival advantage over AZA monotherapy. A possible explanation for these results could be that these agents exhibit strong anti-mitotic properties, which might antagonize incorporation of HMAs into tumor DNA^67,68^. In immune sufficient AML patients, an added benefit of TAK-981 combination therapy may be its potential to activate anti-tumor T and NK cells ^55,69^. While TOPORS is currently not druggable directly, in the interim we propose low dose HMA in combination with TAK-981 as a viable therapeutic strategy to be considered for HR-MDS and AML.

## METHODS

### Human specimens

Cord blood units were supplied by Sydney Cord Blood Bank under the approval of the South Eastern Sydney Local Health District (reference 08/190). Mononuclear cells isolated using lymphoprep (ELITech Group, #1114547), and CD34+ cells were enriched using magnetic beads (Miltenyi Biotec, #130-046-702) on an autoMACS Pro Separator (Miltenyi Biotec) according to manufacturer’s instructions.

### Cell culture and drug treatments

Cell cultures were grown in a humidified incubator at 37°C supplemented with 5% CO_2_. All leukemia cell lines were maintained in RPMI 1640 medium (Life Technologies, 11875-093) containing 10%-20% fetal bovine serum (FBS) (Sigma-Aldrich, F9423-500mL) supplemented with 1X GlutaMAX (Gibco, 35050-061) and 100units/mL of penicillin-streptomycin (Gibco, #15140-122). MDSL and TF-1 were additionally supplemented with 25ng/mL of recombinant human IL-3 (Miltenyi Biotec, 130-095-069), and MDS-L with 50nM β-mercaptoethanol (Sigma-Aldrich, #M6250-100mL). MS5 cells were maintained in Gibco’s α modified eagle’s medium (Gibco, #12571063) supplemented with 10% FBS. The identity of all leukemia cell lines was confirmed by STR profiling at the Garvan Institute of Medical Research and routine mycoplasma testing was performed at the Mycoplasma Testing Facility, UNSW Sydney. The MDS-L cell line was a generous gift from Dr. Kaoru Tohyama (Department of Laboratory medicine, Kawasaki Medical School).

Unless otherwise specified, cells were treated *in vitro* with AZA (Selleck, # S1782), DAC (Selleck, #S1200), TAK-981(Selleck, #S8829 [*in vitro* use] or Takeda [*in vivo* use]), Topotecan (Sapphire BioScience cat. A10939-50, Etoposide (Clifford Hallam Healthcare, cat. 1280860), Hydroxyurea (Selleck, #74-S1896) or GSK3685032 (Selleck cat. E1046) at the indicated concentrations daily for 4 days.

### Capture panel Genotyping

Genotyping of MDS-L and PDX AML-5 was performed as previously described using a capture panel of 111 genes relevant to myeloid malignancies^70^.

### Viral transduction and generation of stable cell lines

pLeGO-iG2-Luc was a gift from Dr Kristoffer Weber^32^. pLKO5d.SFFV.SpCas9.p2a.BSD (Cas9-BsD) was used to generate Cas9-expressing cell lines. The following sgRNA sequences were cloned into SGL40C.EF879S.tagRFP657: AACGGCTCCACCACGCTCGG (sgROSA/CONTROL), CCATGGTGCCTGACTAACAG (sgTOPORS-1), GGACAGTTCAACAAGTTCTG (sgTOPORS-2), TAATATTAGTTCCGTCACAG (sgUBE2K-1), GCAATGACAATAATACCGTG (sgUBE2K-2), ACAGGTTTATCATGACAGTG (sgUBXN7-1), TCAGGTGCAAGTGAAAGTGT (sgUBXN7-2). shRNA lentiviral vectors were purchased from GeneCoiepoeia: GCTTCGCGCCGTAGTCTTA (shCONTROL), GGGCAGAAGATGACTTCAAGG (shTOPORS-1), GCATGATCAGAAGAATCATAG (shTOPORS-2).

Non-replicating lentiviruses were produced in HEK293T cells using the 2^nd^ generation packaging plasmids psPAX2 construct (Addgene #12260), pMD2.G construct (Addgene #12259), and the respective lentiviral transfer plasmid. Lentiviral supernatant supplemented with 8ug/mL polybrene was used for transduction into the respective cell lines.

### ICE Analysis

Genomic DNA was harvested from sgRNA-transduced cells using either the Monarch genomic DNA purification kit (NEB, #T3010S) or QuickExtract DNA extraction solution (Lucigen, #QE0905T) according to manufacturer’s instructions. Genomic regions flanking the expected cut sites for the indicated sgRNAs were PCR amplified using Q5 PCR Master Mix (NEB, #M0541L). PCR products were purified using the Monarch PCR and DNA Cleanup Kit (NEB, #T1030L) prior to Sanger sequencing. Primers for amplification and sequencing were as follows; TOPORS-1: F-TGCCTTCACAGATTAGTCCCCTGG, R-GCCCACTTCTACTCTGAGAACGTG, Seq-TGGAGAGTCAGGCATTTGTGTCTG, TOPORS-2: F-TGCCTTCACAGATTAGTCCCCTGG, R-GCCCACTTCTACTCTGAGAACGTG, Seq-TGCCCTGCTCCTTCATACGAAG, UBXN7-1: F-TGGGAAAGGAGGAGGAATGGGTC, R-CGGGTTCAGGCCATTCTCCTGC, Seq-TGCAATTCTGAAAACAGATCCAGTC, UBXN7-2: F-GCCTCAGCCTCCCAAGGTGTTG, R-GCAGAGCACCACCACACACTCC, Seq-GGCAATGGATAGCTCCTGACAACAC, UBE2K-1: F-CTGCACCCTGCCTCACATGAAG, R-TGTGCTCAATTAACACAACCTGC, Seq-ACACCCCTTCTTTCACCTAGGC, UBE2K-2: F-CCAGCACATTGGGAGGCCAAGG, R GCAGGGAGGGATCATCACTGAAAGG, Seq-AGAGCCAGACTCCGTCTCAGGG. Sequencing traces (.ab1 format) for gene-edited and corresponding wild-type were uploaded to Synthego’s inference of CRISPR edits server for indel analysis (https://ice.synthego.com/#/).

### Dropout Screen

Cas9-expressing MDS-L cells were infected with lentivirus encoding the human Brunello CRISPR knockout pooled library (concentrated lentiviral aliquots were purchased from the Victorian Centre for Functional Genomics, generated from Addgene# 73178) in the presence of 8µg/mL polybrene at a multiplicity of infection of ∼0.3. After 72h, library transduced MDS-L-Cas9 cells were selected with 1μg/mL puromycin for 5 days then left to recover for 72 hours to allow for maximum gene editing. At time-0, ∼5×10^7^ live cells were harvested per replicate and cell pellets underwent same-day nuclei preparation using Qiagen Blood & Cell Culture DNA Maxi Kit (Qiagen, #13362) according to the manufacturer’s instructions. 4.5×10^6^ cells were also harvested and stained for CellTrace staining according to the manufacturer’s instructions to track cellular proliferation in parallel throughout the screen. For dropout screening, Brunello-transduced MDS-L-Cas9 cells were split into 2 treatment arms: AZA (0.3μM) or vehicle (DMSO, 0.000003%). AZA or vehicle was refreshed daily, and cells were passaged every 3-4 days. A minimum of 3.9×10^7^ cells for each replicate and condition were maintained at every passage to preserve library representation. Endpoint cell samples were harvested when the AZA-treated arm underwent 12 cellular divisions as determined by CellTrace staining (alternate colored at each passage). Approximately 6-15×10^7^ cells were harvested for each endpoint samples. sgRNA cassettes were PCR amplified across multiple 50uL reactions, each containing 5ug gDNA, using staggered P7 primers and indexed P5 primers, and 2X NEBNext Q5 high fidelity master mix (NEB, #M0541L) as previously described^25,71^. PCR products were pooled, purified using AMPure XP beads, and 1.0pM of the pooled CRISPR libraries were sequenced on the NextSeq 500 platform using the NextSeq 550 high-output kit with a read length of 1 x 75bp and a 20% PhiX spike-in.

The MAGeCKFlute pipeline was used for processing CRISPR screen data, quality control, and hit identification^26^. Briefly, CRISPR screen data was pre-processed using MAgeCK’s (Version 0.5.9.2) count function with standard parameters to generate sgRNA count tables. Hit identification was performed using MAGeCK MLE (Version 0.5.3) using standard parameters using a 3-condition design (day 0, drug treatment, and DMSO treatment). Functional analysis and visualization of the MAGeCK MLE results were performed using MAGeCkFlute (Version 1.14.0) with standard parameters and normalized with the cell cycle parameter. ClueGO (Version 2.5.9) was used to generate dropout hit cytoscape plots using integrated gene ontology, KEGG, and WikiPathway terms.

### CD34 editing and coculture

Cord blood-derived HSPCs were gene-edited for *TOPORS* or control (sgROSA) using the combined sgRNA-lentiviral and Cas9-mRNA electroporation approach as previously described^34^.

For transduction, lentiviral supernatants were loaded in RetroNectin-coated plates then centrifuged at 1000 x g for 90mins at room temperature. After washing plates, 2mL of CD34^+^ cells were added at a density of 1-2×10^5^ cells/mL and incubated for 72 hours in a 37°C incubator. Transduced cells (tagRFP657^+^) were sorted for CD34^+^ using anti-human CD34-PE antibody (BD Biosciences, # 555822) on the FACSAriaIII (Becton Dickinson), then electroporated with GFP-Cas9 mRNA (Dharmacon, #CAS11860) using a Neon electroporator (Invitrogen). Electroporated cells were immediately transferred to 0.5ml pre-warmed media in a 24-well plate and returned to the 37°C incubator. Transfection efficiency was checked by GFP expression 16-24 hours after electroporation and gene editing efficiencies were checked 4-5 days after transfection. Efficient editing of the *TOPORS* locus was confirmed through ICE analysis from approximately 10000 cells per sample.

24 hours after combinatorial trannsduction and electroporation was performed, 10000 TOPORS-edited or control CD34^+^ cord blood-derived HSPCs were seeded into 24-well plates in an MS5 co-culture system. Briefly, MS5 cells were plated into 24-well plates with 1.5×10^5^ cells/well and left to form a confluent monolayer overnight. CD34^+^ cells were then plated onto MS5 monolayer in MyeloCult H5100 media (Stemcell Technologies, #05150) and cells treated with 0.5µM AZA for 4 consecutive days. On Day 5, cells were collected from culture plates and sorted for human CD34^+^ cells. MS5 cells were excluded using anti-mouse CD105-eFluor450 antibody (Invitrogen, #48-1051-82).

### Colony forming assay

1000 sorted CD34^+^ cells, or 1000 MDS-L cells were seeded per 1mL of MethoCult H4434 (StemCell Technologies, #04434) in 35mm dishes. Healthy colonies were scored after 10-14 days according to manufacturer’s instructions. MDS-L primarily produce CFU-GM. Only colonies greater than 40 cells were scored.

### Competitive proliferation assay

Cas9-expressing MDS-L cells were transduced with sgRNA/tagRFP657^+^ lentiviral constructs and were left unsorted for tagRFP. Cells were treated daily with 0.3µM AZA or DMSO for 16 days. The proportion of tagRFP657^+^ cells were measured using the BD LSRFortessa SORP at pre-treatment levels and every 4 days up until day 16.

### EC50 measurements

Cells seeded in 96-well plates were exposed to inhibitors at indicated concentrations. Following drug treatment, DAPI was added to each well to a final concentration of 0.25µg/mL. Plates were analyzed on the Attune NxT (Invitrogen) using the 96-well sampler with fixed volume analysis to record the absolute number of DAPI^-^ and fluorescent protein positive cells (tagRFP657^+^ for sgRNA-transduced samples; GFP^+^ for shRNA-transduced samples). Dose-response curves were fitted using the log(inhibitor) vs normalized response 4-parameter variable slope in GraphPad Prism (v10.1.1).

To test drug synergy, cells were treated with inhibitor in a 2D dose matrix, and viability determined by MTS assays (Promega, #G5430). Synergy scores were calculated using https://synergyfinder.fimm.fi/. ZIP scores less than -10 indicated an antagonistic interaction between both drugs, from -10 to 10 indicated an additive interaction, and greater than 10 indicated a synergistic interaction between both drugs.

### Apoptosis analysis using annexin V/Propidium Iodide (PI)

Apoptosis analysis by Annexin V/PI staining was performed using Abcam’s Annexin V-FITC kit (Abcam, #ab14085) according to the manufacturer’s instructions.

### γH2AX and cell cycle flow cytometry

Cells were washed, fixed with 4% formaldehyde in PBS for 10 minutes at room temperature, permeabilized with ice-cold 90% v/v methanol and incubated for 5 minutes at room temperature, then resuspended in FACs buffer (2% FBS, 1mM EDTA in PBS); all in the dark or low lighting.

For γH2AX staining, cells were blocked using human Fc Block (BD, #564220) then stained with γH2AX antibody (2µg/mL, Abcam, #ab26350) for 1 hour at room temperature, followed by anti-mouse DyLight 488 antibody (1:500 in 0.1% NP40/PBS, Abcam, #ab96879) for 30 minutes at room temperature; all in the dark or low lighting. Cells were analysed on the LSRFortessa SORP.

For cell cycle analysis, fixed cells stained with DAPI (1µg/mL, BD Biosciences, #564907) were analyzed on the LSRfortessa SORP with V450 and UV450 channels set to linear. Samples were run at low speed and voltages adjusted until the G1 peak sat as close to 100 as possible. Data was analyzed on FlowJo (Version 10.7.1) applying the Dean Jett Fox pragmatic model.

### Comet Assay

Cell pellets were washed once with ice-cold PBS (without Mg2+ and Ca2+), then resuspended at 1×10^5^ cells/mL in ice-cold PBS and analysed using the comet assay kit (Abcam, #ab238544) according to manufacturer’s instructions with gel electrophoresis run for 20 minutes at 3 volts/cm.

### RNA Seq and data analysis

Total RNA was extracted using the Bioline Isolate II RNA extraction Kit (Bioline, BIO-52072) according to the manufacturer’s instructions. Residual DNA was eliminated by using the RNase-free DNase Qiagen Kit (Qiagen, #79254). RNA-seq libraries were prepared using the Illumina Stranded mRNA prep Ligation kit performed as per manufacturer’s instructions and libraries sequenced on the NovaSeq 6000 platform using 1 lane of a 2 x 100bp SP flow cell.

Raw FASTQ files assessed for quality control through FastQC and sequencing adaptors and low quality reads removed using BBmap. Paired-end reads were mapped to the hg38 reference genome using STAR (Version 2.7.0) and quantified using featureCounts (Version 2.0.0). The parameters “--outFilterMismatchNoverLmax” and “--alignEndsType” of STAR (Version 2.7.0) aligner was set to “0.05” and “EndToEnd” to filter out reads harbouring artifact mismatches from the mapping process. Read count matrices were generated with FeatureCounts using default arguments with “requireBothEndsMapped” and “countChimericFragments” options.

rMATS (Version 4.1.2) was used to detecting splicing events and differentially splicing^49^. Spliced events were filtered using an FDR cut-off of less than 0.05 and inclusion level difference greater than |0.1|. Five types of splicing events were captured: exon skipping, Intron retention, alternative 3’ splice site, alternative 5’ splice site, and mutually exclusive exons.

Unsupervised hierarchical clustering analysis was used to assess the power of differentially spliced events in separating samples of different comparisons aligned with their labels (MRAZA, MRDMSO, MT1AZA and MT1DMSO). First z-score normalization was performed on processed Inclusion Level table to adjust for the signal to noise ratio. The K-means algorithm was used to perform unsupervised hierarchical clustering on both genes and labels. The number of K for labels was set to four. To obtain a consensus k-means clustering, 100 k-means runs were executed for both labels and genes simultaneously.

To identify differentially expressed genes, DESeq2 R package (Version 1.34.0) was used. First gene count tables were normalized and genes with normalized read counts less than 10 were filtered out. RNA binding factor motif density scanning was performed using the RMAPs2 server (http://rmaps.cecsresearch.org/#about-section)^50^. Raw outputs from rMAPS were uploaded onto the server and analyzed for RNA binding factor motif densities using default settings. Overrepresentation and transcription factor regulatory target analyses were performed using the MetaScape server (https://metascape.org/gp/index.html). Filtered lists (FDR< 0.05, |Log_2_Fold-Change|>1) were uploaded for express analysis.

### Quantitative RT-PCR

RNA was reverse transcribed into cDNA using the QuantiTect reverse transcription kit (Qiagen, #205311) as per manufacturer’s instructions. cDNA was amplified PowerUP SYBR green master mix (Applied Biosystems, #A25776) on a BioRad CFX96 real-time PCR machine using the following primers; β-Actin: F AGCACTGTGTTGGCGTACAG, R AGAGCTACGAGCTGCCTGAC,PSMB4: F GATCCGGCGTCTGCACTTTACAG, R CATGTCTGCGGCAATCACCACTC,TOPORS: F GACACCGACCTAGCTTTCTGGG, R TTTGCTAGTGCCAGCTTTAGGTG. Relative mRNA expression levels of *TOPORS* was normalized to either β-Actin or PSMB4 using the 2^−ΔΔ*Ct*^ method.

### End-specific qPCR for HpaII-digestable LINE-1 promoters

The LINE-1 end-specific qPCR (ESPCR) to quantify demethylated LINE-1 promoters was performed as previously described^72^ with minor modifications. Primers, facilitator oligonucleotides (Foligos), and probes were synthesized according to the published sequences^72^, except for the HpaII-cut specific reverse Foligo: 5’-TGGCTGTGGGTGGTGGGCCTCGTAGAGGCCCTTTTTTGGTCGGTACCTCAGATGGAAATGTCTT/3ddC/-3’. For a 20 μL reaction, 10 μL of master mix was added to 10 μL of DNA. The master mix composition included 4.0 μL of 5X ESPCR buffer, 1.2 μL of 50 mM MgCl2, 1.0 μL of 20× DraI-cut specific oligonucleotide mixture, 1.0 μL of 20× HpaII-cut specific oligonucleotide mixture, 0.2 μL of HpaII enzyme (NEB, #R0171L), 0.05 μL of DraI enzyme (NEB, #R0129L), 0.1 μL of Hot Start Taq DNA Polymerase (NEB, #M0495L), and 2.45 μL of nuclease-free water. qPCR was conducted using a BioRad CFX96 real-time PCR machine under the following conditions: 37°C for 15 min; 90°C for 5 s; 95°C for 2 min; then 10 cycles of 90°C for 5 s, 95°C for 15 s, 60°C for 1 min, and 68°C for 20 s; followed by 40 cycles of 95°C for 15 s, 65°C for 40 s, and 68°C for 20 s. FAM and HEX readings were recorded at 65°C during the final stage of 40 cycles. The efficiency of the FAM and HEX reactions was calculated using a standard curve generated from 4 pg to 4 ng of DNA from AZA-treated RKO cells. The relative demethylation of LINE-1 promoters in samples was calculated using ΔΔC_T_, normalised to the average vehicle ΔΔC_T_.

### Western Blotting

Nuclei protein from MOLM-13-Cas9 cells transduced with either sgROSA or sgTOPORS-1 treated with 2 doses of DAC or vehicle over 24hrs was extracted in RIPA buffer supplemented with protease inhibitor (Roche, #04693159001). 10ug of protein was boiled in NuPAGE LDS sample buffer (Invitrogen, #NP0007) containing 0.1M DTT and samples run on 4-12% Bis-Tris gel. (Invitrogen, #NP0323) and transferred to PVDF (Thermofisher Scientific, # 88518). Blocked membranes were probed overnight at 4°C with primary antibodies diluted 1:1000 in 5% skim milk in TBST. Primary antibodies used were: anti-DNMT1(Cell Signaling Technology #5032) or anti-SUMO-2/3 (MBL Life Science #M114-3). Membranes were stripped in 15g glycine, 1g SDS, 10ml Tween-20, pH2.2 then re-probed with re-probed with anti-β actin (Santa Cruz Biotechnology, #sc-47778). HRP-conjugated anti-rabbit and anti-mouse secondary antibodies (Dako #P0448 and #P0260) were diluted 1:2000 and incubated for one hour at room temperature. Chemiluminescent signal was detected using Clarity Western ECL Substrate (Biorad, #1705061) and visualised using the iBright CL1500 Imaging System (Invitrogen).

### AZA-MS

AZA-MS was used to measure levels of 5 aza-dC and 5 me-dC in genomic DNA as previously described ^38^.

### Nuclear Proteomics

Nuclear lysates were prepared by resuspending cell pellets in lysis buffer (10mM Tris pH 8.0, 10mM NaCl, 0.2% NP40) supplemented with protease inhibitors (Roche, # 4693159001) and phosSTOP (Roche, # 4906837001), incubating for 10 min on ice, and pelleting nuclei at 1450g. The nuclear pellet was lysed in ice-cold RIPA (50mM TrisHCl pH7.5, 150mM NaCl, 1% Triton X-100, 0.5% Deoxycholic acid, 0.1% SDS) supplemented with protease inhibitors and phosSTOP, with Benzonase added to a final concentration of 50units/mL. Lysates were sonicated on the Bioruptor (Diagenode) at 30s on and 30s off for a total of 3 cycles and incubated on ice for 15 minutes. Samples were spun down at 8000 x g for 15 minutes at 4°C and the protein supernatant was collected into a fresh tube. Quantified nuclear protein lysate was used for mass spectrometry-based label-free protein quantification as described^73^.

LC-MS/MS raw files were pre-processed and analyzed using the MaxQuant software suite (Version 1.6.2.10.43) for feature detection, protein identification and quantification, and sequence database searches were performed using the integrated Andromeda search engine as previously described^73^. MaxQuant pre-processed files were loaded into LFQ-Analyst^74^ for label-free protein quantification using cutoffs of FDR< 0.05, |Log_2_Fold-Change|>1, with a perseus-type imputation, Benjamini Hochberg FDR correction, and inclusion of single peptide identifications.

### LC-MS/MS of Ni-NTA enrichment of 10His-SUMO1ylated proteins

A human codon optimised sequence encoding HHHHHHHHHH-MSDQEAKPSTEDLGDKKEGEYIKLKVIGQDSSEIHFKVKMTTHLKKLKESYCQRQGVPMNSLRFLFEGQRIADNHTPKELGMEEEDVIEVYQEQTGG (10His-SUMO1) was fused to DasherGFP separated by the T2A peptide EGRGSLLTCGDVEENPGP and cloned into lentiviral vector pD2109-EF1 by Atum Bio (Fig. S6A), and lentiviral particles used to stably transduce gene-edited MOLM-13 cells. DasherGFP^+^tagRFP^+^ gene-edited MOLM-13 cells purified by FACS were cultured in quadruplicate and treated with 41nM DAC or vehicle daily for 3 days. 10His-SUMO1ylated proteins were enriched from 6M guanidine whole cell extracts using Ni-NTA beads following the detailed procedure of^54^, stopping the enrichment process at Step 47. Proteomic data was recovered from the enriched proteins using LC-MS/MS of trypsin digests as for nuclear proteins from gene edited MDS-L cells.

### Xenotransplantation into MISTRG mice

MISTRG mice^33^ were bred and maintained in SPF conditions in HEPA filtered cages at ≤5 mice per cage under protocols approved by the Animal Care and Ethics Committee of the University of New South Wales.

### Cord blood CD34+ cells

MISTRG neonates (2–3 days old) were xeno-transplanted by intrahepatic injection with 2000–3000 CD34+ human cord blood cells in 25µL neutral saline as described ^33^, with the omission of neonatal irradiation. Engraftment levels were quantified by flow cytometry of tail bleeds performed no earlier than 7 weeks of age and expressed as the percentage of human CD45+ (huCD45+) cells to total huCD45+ plus mouse CD45+ (moCD45+) cells in the tissue sample. To randomly distribute treatments across cages, mice were rank randomized according to sex and %huCD45+ in cohorts equal to the number of treatment groups prior to commencement of drug treatments, which then proceeded as described in the Figures.

### MOLM-13 cells

Warmed adult MISTRG were tail vein injected with 5×10^4^ (females) or 10^5^ (males) luciferase^+^ MOLM-13 cells i.v. in an inoculum of 0.1mL neutral saline. Injected mice were maintained with a constant supply of sterile filtered 0.27mg/mL enrofloxacin (Bayer) antibiotic in their drinking water. Engraftment was first assessed about 10 days after inoculation using an IVIS SpectrumCT Preclinical In Vivo Imaging System (PerkinElmer) for whole body luminescence imaging under isoflurane anaesthesia, as described^75^. Mice were then rank randomized according to sex and pre-treatment luminescence flux in cohorts equal to the number of treatment groups prior to commencement of drug treatments.

### Primary AML cells

The origin and maintenance of patient AML-5 cells as continuous mouse xenografts in NSG strain mice is described^56^. Freshly thawed AML-5 xenograft mouse bone marrow and spleen cells (>95% huCD45+) were injected into 8 week old MISTRG males and females (1.25×10^6^ cells per mouse) that had received 250 cGy sub-lethal irradiation 24h prior. Xeno-engrafted mice were ranked randomized according to sex and weight in cohorts equal to the number of treatment groups. Drug treatment cycles then commenced 11 days after inoculation and proceeded as described in Figures. Engraftment levels were quantified by flow cytometry of tail bleeds as above, sampled at ∼1 week intervals.

### Xenograft drug treatments

HMA stocks (30mg/mL for AZA; 10mg/mL for DAC) dissolved in anhydrous DMSO and stored at -80C under argon in small aliquots were diluted into neutral saline immediately before use and administered i.p. or s.c. at the concentrations specified in a bolus of 5mL/kg. TAK-981prepared as a 4-5mg/mL working solution in 20% hydroxypropyl β-cyclo-dextrin vehicle (HPBCD; Onbio Inc.) at pH 3.5–4 (stored at -80C for ≤8 weeks), was diluted in HPBCD vehicle if necessary and injected i.p. at 5mL/kg. Tetrahydrouridine (THU, Abcam) dissolved to 2mg/mL in neutral saline (stored at -20C for ≤12 weeks) was injected i.p. at 5mL/kg.

### Data Availability

Data from the CRISPR dropout screen and RNA sequencing have been deposited at GEO under accession #GSE261339. The mass spectrometry proteomics data have been deposited to the ProteomeXchange Consortium with the dataset identifier PXD050539.

## ACKNOWLEDGMENTS

The authors thank the staff and donors of the Sydney Cord Blood Bank for providing cord bloods for research, Takeda Pharmeceuticals for provision of TAK-981 for use in mice, Prof Kaylene Simpson (Peter MacCallum Cancer Centre) for provision and advice with the CRISPR lentiviral library, Forrest Koch (UNSW) for assistance with genotyping, Dr Ivo A. Hendriks (University of Copenhagen) for advice on SUMO pulldown assays, Dr Antony Rongvaux (Fred Hutchinson Cancer Research Centre) for advice on MISTRG colony maintenance and xenotransplantation, Dr Gil Prive and Dr. Brian Raught (University of Toronto) for stimulating discussions. Some of the data presented in this work were acquired by personnel and/or instruments of the National Imaging Facility, a National Collaborative Research Infrastructure Strategy (NCRIS) capability, at the Mark Wainwright Analytical Centre (MWAC) of UNSW Sydney, which is funded in part by the Research Infrastructure Programme of UNSW. The Victorian Centre for Functional Genomics is funded by the Australian Cancer Research Foundation, Phenomics Australia through funding from the Australian Government’s National Collaborative Research Infrastructure Strategy program, and the Peter MacCallum Cancer Centre Foundation. This work was supported by Postgraduate Awards from UNSW Sydney (P.T, A.A, X.Z, G.S.B, E.G) and Translational Cancer Research Network-a Translational Cancer Research Centre funded by the Cancer Institute NSW (P.T.); the Anthony Rothe Memorial Trust (J.T, J.E.P, P.C); Cancer Council NSW (RG 23-08, P.C), Ideas Grant from the National Health and Medical Research Council of Australia (GNT2011627; C.J.J, J.E.P); Research Fellowship from the National Health and Medical Research Council of Australia (APP1157871; R.B.L.); program grants from the Leukemia Lymphoma Society (LLS)-Snowdome Foundation-Leukaemia Foundation (6620-21; J.E.P), Cancer Institute NSW (TPG2152; J.E.P, J.T) and Medical Research Future Fund (MRF1200271; J.E.P). Children’s Cancer Institute Australia is affiliated with UNSW Sydney and The Sydney Children’s Hospitals Network.

## AUTHOR CONTRIBUTIONS

P.T, S.S, S.J, Md. I. I, L. Z, M.J.R, A.A, M.N, X.Z, G.S.B, C.H.S, E.G, O.S, S.M, C.E.T, P.C, P.M.K, S.K.B, K.A.M, J.A.I.T, C.J.J performed the research and analyzed the data; M.R, H. A-R, J.L, P.C., K.A.M, R.B.L and C.W provided key reagents, discussed, and interpreted the data. The study was conceived by J.E.P. The manuscript was written by P.T. and C.J.J with contributions from J.A.I.T and J.E.P.

