## Supplemental Figures and Legends for "Genome-Wide CRISPR-Cas9 Screening Identifies a Synergy between Hypomethylating Agents and SUMOylation Blockade in MDS/AML"

**Genome-Wide CRISPR-Cas9 Screening Identifies a Synergy between Hypomethylating Agents and SUMOylation Blockade in Myelodysplastic Syndromes and Acute Myeloid Leukemia**

Peter Truong et al

### Supplemental Table Legends

#### Table S1. Genotypes

Myeloid capture panel genotyping of MDS-L cells and patient derived xenograft used in this work.

#### Table S2. sgRNA counts and β scores

(A) Raw sgRNA counts and (B) corresponding β Scores for AZA and vehicle (DMSO) treatment, determined by MAGeCKFlute analysis of the MDS-L genome-wide CRISPR/Cas9 dropout screen shown in Fig. 1.

#### Table S3. EC50 values for sgCONTROL and sgTOPORS cells

EC50 values (Prism 10.1.1; log[drug] vs. survival – variable slope, 4 parameters) calculated from data shown in Fig. 7.

#### Table S4. EC50 values for AZA/DAC/TAK981 combinations

EC50 values (Prism 10.1.1; log[drug] vs. survival – variable slope, 4 parameters) calculated from data shown in Fig. 9B–C.

### Supplemental Figures and Legends

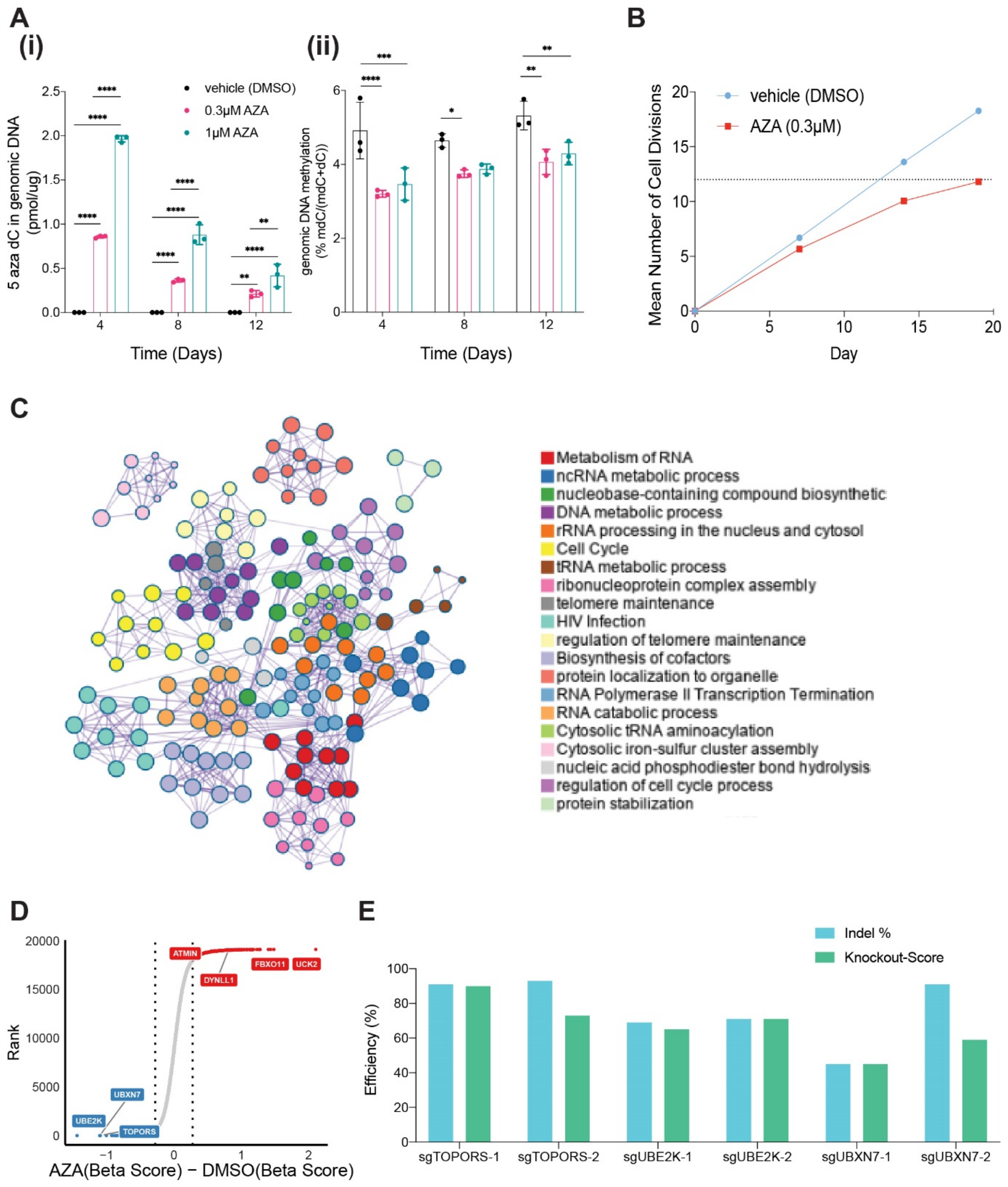


**Figure S1**

**Figure S1. Genome-wide CRISPR-Cas9 dropout screening identifies novel genetic determinants of AZA-sensitivity**

(A) (i) Incorporation of 5 aza-dC, and (ii) methylation at dC, in genomic DNA (both determined by LC-MS) in MDS-L cells exposed to AZA daily for 4 days followed by an 8-day drug holiday. Shown are mean ± SD and summary of 2-way ANOVA: non-significant differences not shown, * $P \leq 0.05$, ** $P \leq 0.01$, *** $P \leq 0.001$, **** $P \leq 0.0001$. (B) Number of cell divisions since start of drug selection, determined by cycling between CellTrace-Yellow and -Far Red staining in a sub-culture tracked alongside the genome-wide CRISPR-Cas9 screen culture. (C) Cytoscape plot highlighting cell-essential modules of biological processes enriched in hits that dropped out from both AZA and DMSO treatment arms. (D) All gene targets recovered in the CRISPR-Cas9 screen ranked according to their differential β score calculated by subtracting the vehicle (DMSO) β score from the AZA β score. (E) Efficiency of Indel generation and the KO score generated by specific Indel distributions, as determined by applying the "Inference of CRISPR Edits" (ICE) algorithm to data generated via Sanger sequencing of specific sgRNA binding site flanks in genomic DNA pools.

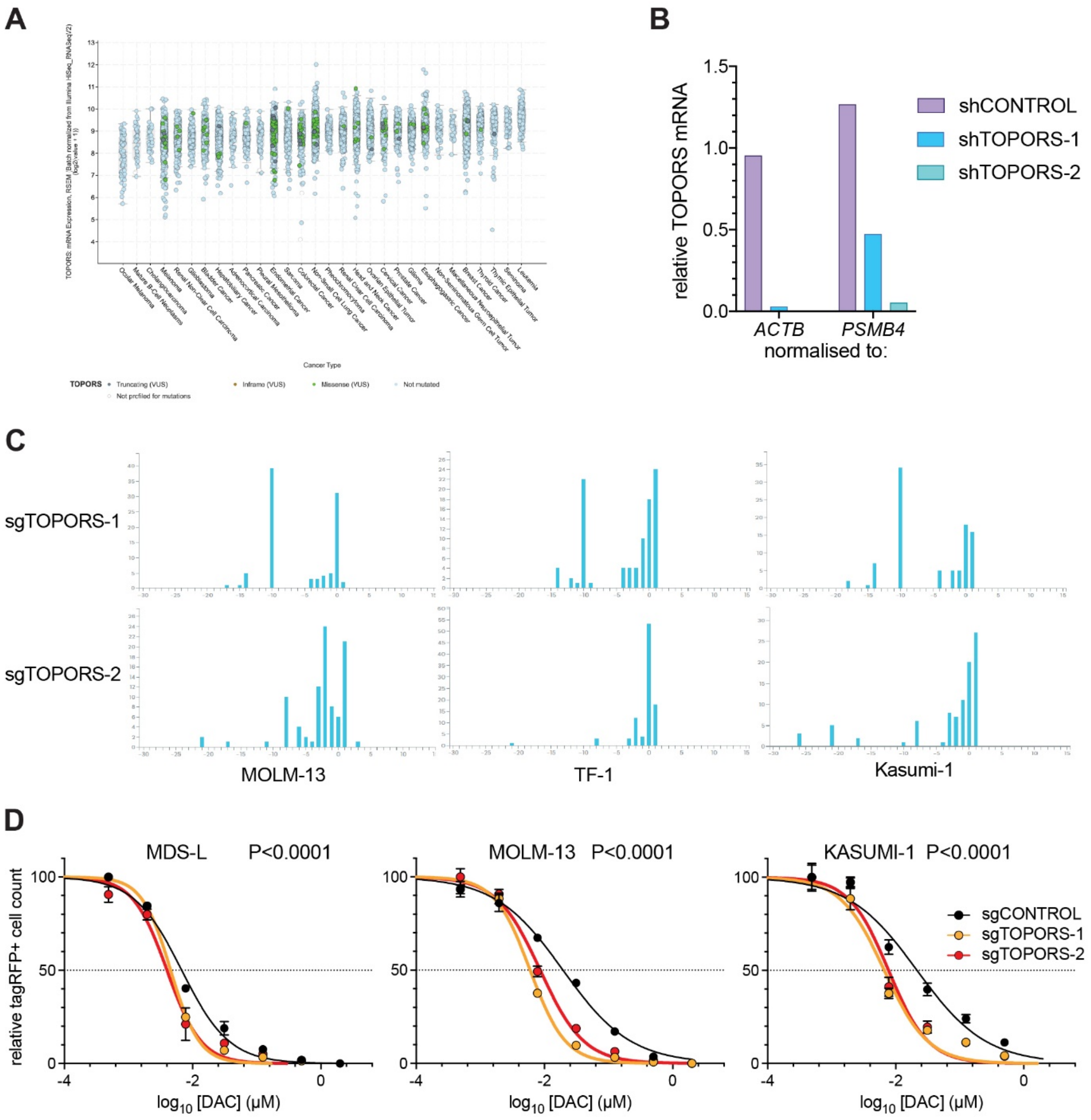
A
Cancer Type
TOPORS
Truncating (VUS)
Inframe (VUS)
Missense (VUS)
Not mutated
Not profiled for mutations
B
relative TOPORS mRNA
1.5
1.0
0.5
0.0
ACTB
PSMB4
normalised to:
shCONTROL
shTOPORS-1
shTOPORS-2
C
sgTOPORS-1
sgTOPORS-2
MOLM-13
TF-1
Kasumi-1
D
MDS-L P<0.0001
MOLM-13 P<0.0001
KASUMI-1 P<0.0001
relative tagRFP+ cell count
100
50
0
-4
-2
0
log10 [DAC] (μM)
sgCONTROL
sgTOPORS-1
sgTOPORS-2

**Figure S2**

**Figure S2. Loss of TOPORS sensitizes MDS and AML cell lines to AZA.**

(A) mRNA expression (RSEM normalized expression values log2(value+1)) and mutational profile of *TOPORS* across cancer types using the TCGA dataset (n=10,071) annotated from the cBioPortal database. (B) Relative mRNA levels for *TOPORS* in cells expressing single shRNAs as determined by qPCR normalized to housekeeping genes *ACTB* (β-Actin) or *PSMB4*. (C) Indel distributions in the *TOPORS* gene, as determined by ICE, in Cas9+ AML cell lines transduced to express the indicated sgRNAs. (D) Dose-survival plots of normalised tagRFP+ cell counts following 4 days of daily treatment with the indicated DAC concentrations of AML cell lines expressing Cas9 plus single sgRNAs or a non-targeting control sgRNA. P-values are from extra sum-of-squares F tests.

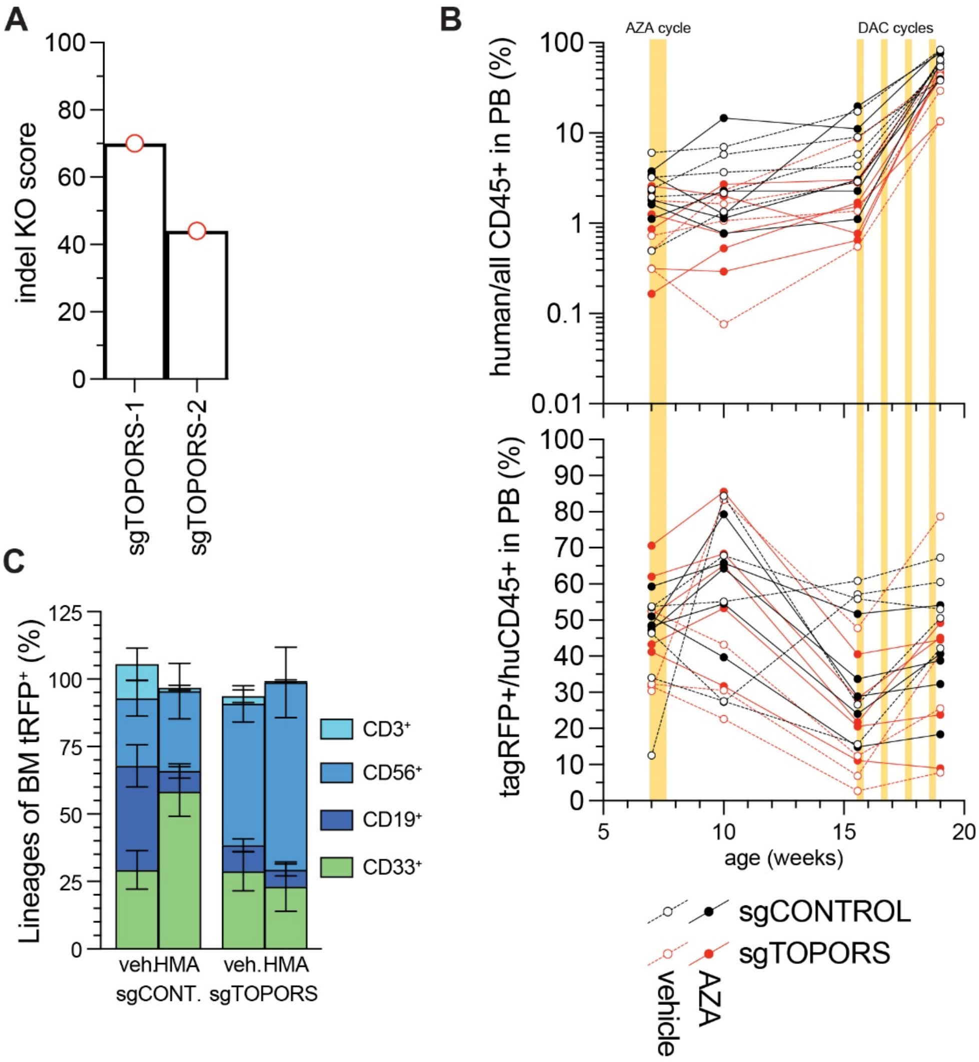
A
100
80
60
40
20
0
indel KO score
sgTOPORS-1
sgTOPORS-2
B
AZA cycle
DAC cycles
100
10
1
0.1
0.01
human/all CD45+ in PB (%)
100
90
80
70
60
50
40
30
20
10
0
tagRFP+/huCD45+ in PB (%)
5
10
15
20
age (weeks)
sgCONTROL
sgTOPORS
vehicle
AZA
C
125
100
75
50
25
0
Lineages of BM tRFP⁺ (%)
CD3⁺
CD56⁺
CD19⁺
CD33⁺
veh.HMA
veh.HMA
sgCONT.
sgTOPORS

Figure S3

**Figure S3. Targeting TOPORS spares healthy hematopoiesis.**

(A) Indel KO scores generated by ICE for CD34+ cells seeded into colony assays in Fig. 2B. (B) Tracking of (top) blood % huCD45+moCD45- cells amongst all CD45+ cells and (bottom) blood % tagRFP+ amongst huCD45+moCD45- cells for each engrafted mouse across all sampling points. Drug treatment days are indicated by bars. (C) Distributions between lineages of bone marrow tagRFP+CD34-huCD45+moCD45- cells at end point ± SEM; n = 4–5.

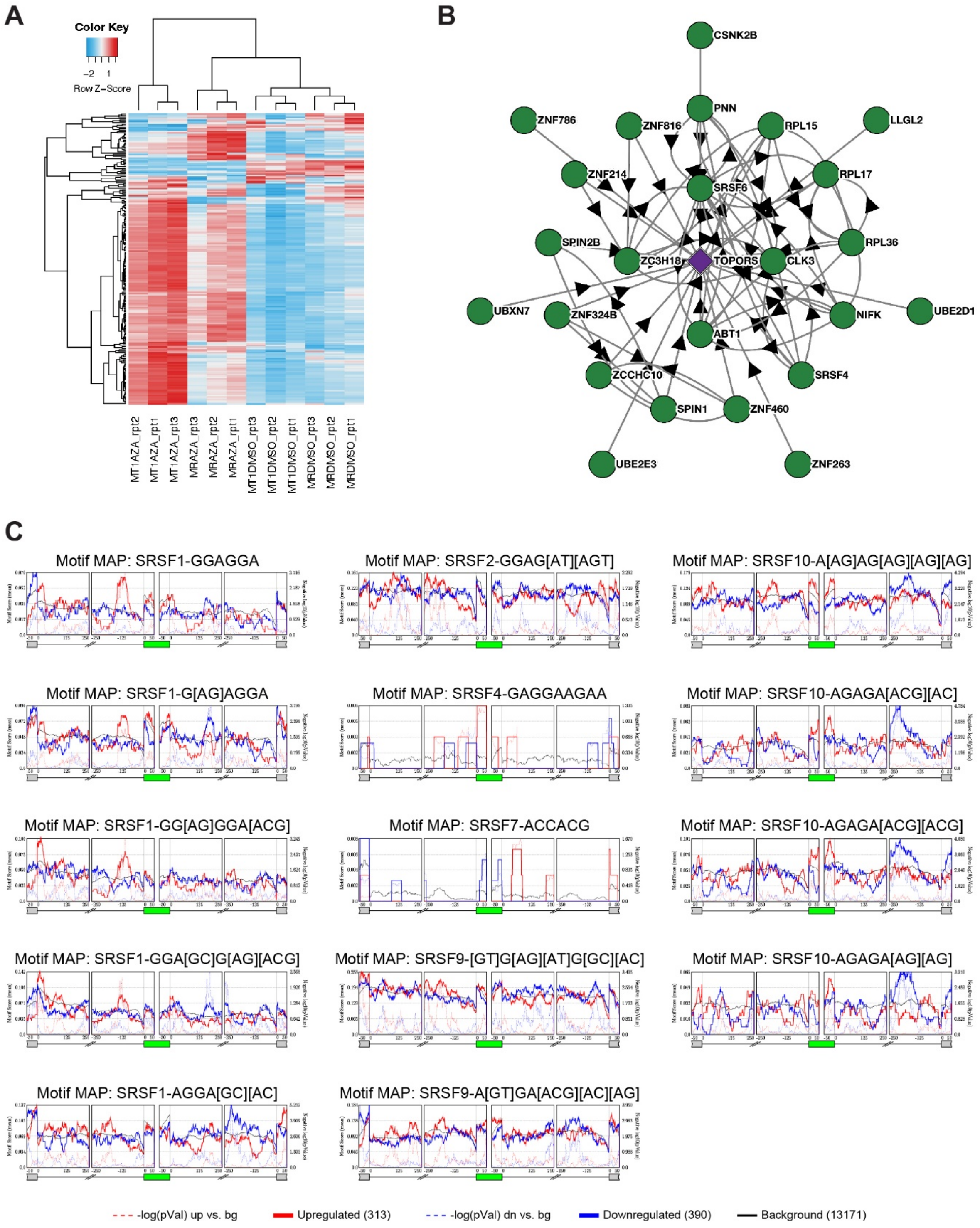
A
Color Key
Row Z−Score
MT1AZA_rpt2
MT1AZA_rpt1
MT1AZA_rpt3
MRAZA_rpt3
MRAZA_rpt2
MRAZA_rpt1
MT1DMSO_rpt3
MT1DMSO_rpt2
MT1DMSO_rpt1
MRDMSO_rpt3
MRDMSO_rpt2
MRDMSO_rpt1
B
CSNK2B
PNN
ZNF786
ZNF816
RPL15
LLGL2
ZNF214
RPL17
SRSF6
SPIN2B
RPL36
ZC3H18
TOPORS
CLK3
UBXN7
ZNF324B
NIFK
UBE2D1
ABT1
ZCCHC10
SRSF4
SPIN1
ZNF460
UBE2E3
ZNF263
C
Motif MAP: SRSF1-GGAGGA
Motif MAP: SRSF2-GGAG[AT][AGT]
Motif MAP: SRSF10-A[AG]AG[AG][AG][AG]
Motif MAP: SRSF1-G[AG]AGGA
Motif MAP: SRSF4-GAGGAAGAA
Motif MAP: SRSF10-AGAGA[ACG][AC]
Motif MAP: SRSF1-GG[AG]GGA[ACG]
Motif MAP: SRSF7-ACCACG
Motif MAP: SRSF10-AGAGA[ACG][ACG]
Motif MAP: SRSF1-GGA[GC]G[AG][ACG]
Motif MAP: SRSF9-[GT]G[AG][AT]G[GC][AC]
Motif MAP: SRSF10-AGAGA[AG][AG]
Motif MAP: SRSF1-AGGA[GC][AC]
Motif MAP: SRSF9-A[GT]GA[ACG][AC][AG]
-log(pVal) up vs. bg
Upregulated (313)
-log(pVal) dn vs. bg
Downregulated (390)
Background (13171)

**Figure S4**

**Figure S4. Multi-omic approaches reveal widespread mis-splicing of DDR genes in AZA-treated TOPORS-edited MDS-L cells.**

(A) Heatmap of transcriptomes clustered according to their gene expression level for E2F target genes (n=200). (B) BioPlex interaction map depicting TOPORS interactors. (C) Overrepresentation analyses of skipped exon events detected in AZA-treated *TOPORS*-edited MDS- L cells compared to AZA-treated control cells.

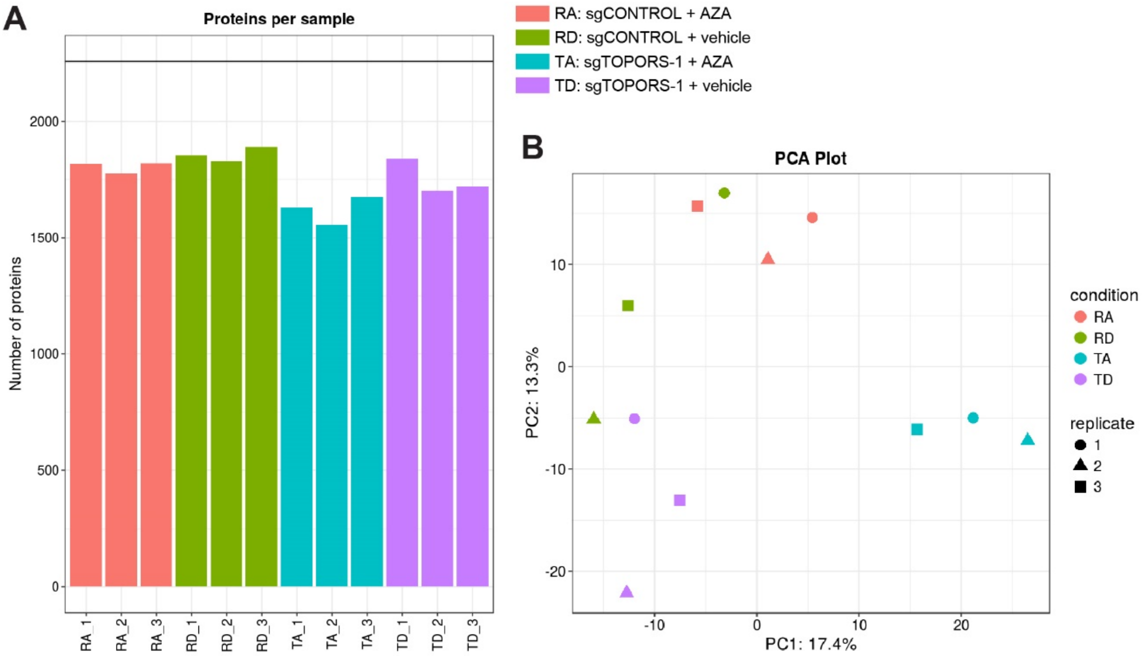
A
Proteins per sample
Number of proteins
2000
1500
1000
500
0
RA_1
RA_2
RA_3
RD_1
RD_2
RD_3
TA_1
TA_2
TA_3
TD_1
TD_2
TD_3
RA: sgCONTROL + AZA
RD: sgCONTROL + vehicle
TA: sgTOPORS-1 + AZA
TD: sgTOPORS-1 + vehicle
B
PCA Plot
PC2: 13.3%
PC1: 17.4%
10
0
-10
-20
20
condition
RA
RD
TA
TD
replicate
1
2
3

**Figure S5**

**Figure S5. Nuclear proteomics reveal depletion of global nucleotide excision repair factors in AZA-treated TOPORS-edited MDS-L cells.**

Summaries of nuclear proteomics data, generated using LFQ Analyst. (A) The number of proteins detected per sample. (B) Principal component analysis of nuclear proteomes.

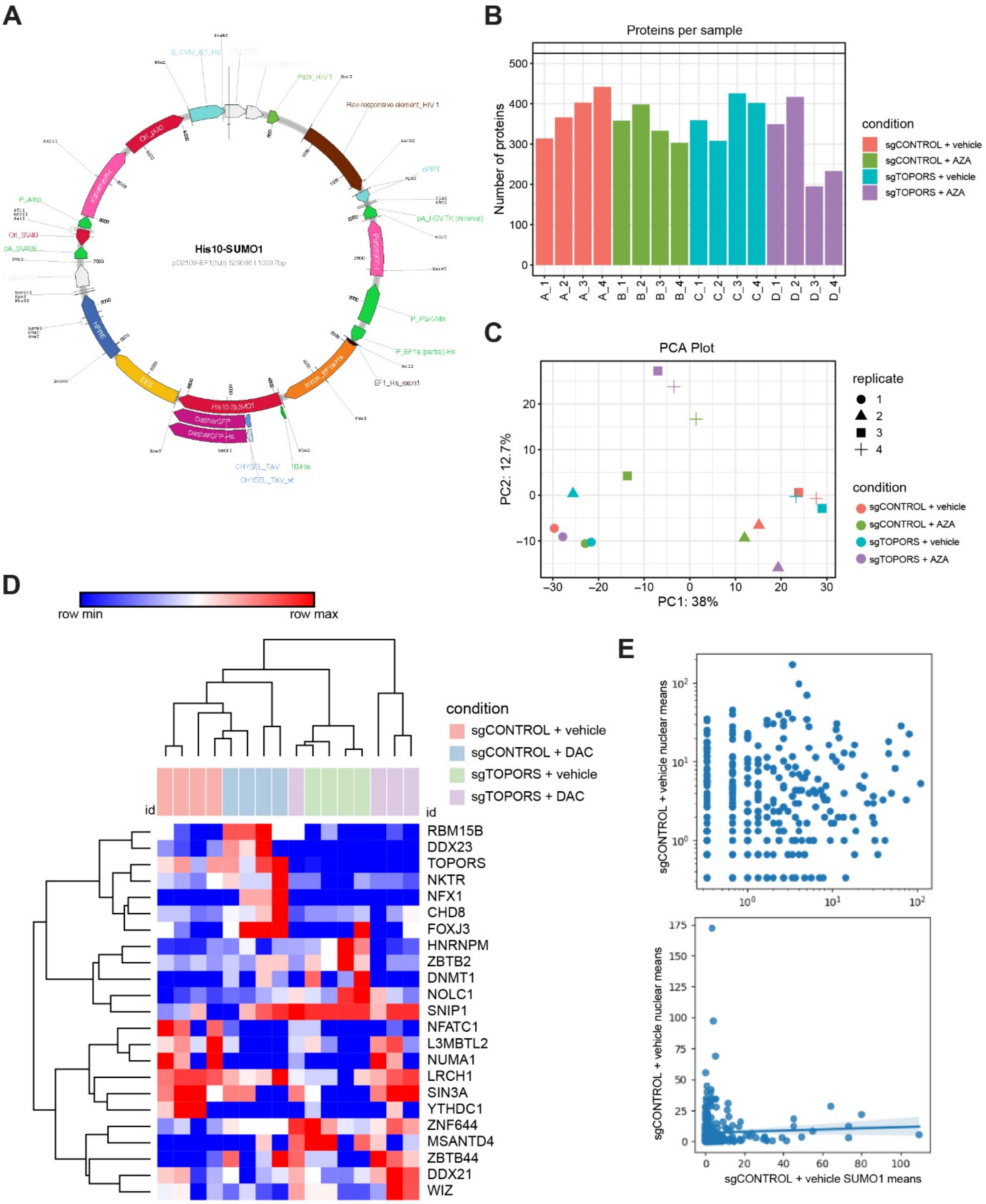
A
His10-SUMO1
B
Proteins per sample
Number of proteins
condition
sgCONTROL + vehicle
sgCONTROL + AZA
sgTOPORS + vehicle
sgTOPORS + AZA
C
PCA Plot
PC1: 38%
PC2: 12.7%
replicate
condition
D
row min
row max
condition
sgCONTROL + vehicle
sgCONTROL + DAC
sgTOPORS + vehicle
sgTOPORS + DAC
RBM15B
DDX23
TOPORS
NKTR
NFX1
CHD8
FOXJ3
HNRNPM
ZBTB2
DNMT1
NOLC1
SNIP1
NFATC1
L3MBTL2
NUMA1
LRCH1
SIN3A
YTHDC1
ZNF644
MSANTD4
ZBTB44
DDX21
WIZ
E
sgCONTROL + vehicle nuclear means
sgCONTROL + vehicle SUMO1 means

**Figure S6**

**Figure S6. TOPORS-editing does not reduce SUMOylation of DNMT1 in DAC-treated AML cells.**

Summaries of Ni-NTA enriched proteomics data. (A) Map for plasmid encoding lentivirus to co-express 10xHis-SUMO1 and DasherGFP separated by a 2A peptide. (B) The number of proteins detected per sample (LFQ Analyst). (C) Principal component analysis of Ni-NTA enriched proteomes (LFQ Analyst). (D) Heatmap of unsupervised hierarchical clustering of 23 proteins that were significantly differentially abundant according to one way ANOVA (Scaffold 5.3.2) across all Ni-NTA pull-downs from quadruplicate cultures of 42 nM DAC/vehicle-treated gene-edited MOLM-13 cells. (E) Correlation plots for Ni-NTA captured compared to nuclear proteomics. x-axes: mean normalised total spectra values (Scaffold 5.3.2) for all proteins detected in MOLM-13/Cas9+sgCONTROL Ni-NTA pull-downs plotted on (top) log scale or (bottom) linear scale. y-axes: mean normalised total spectra values (Scaffold 5.3.2) for the same proteins detected in MDS-L/Cas9+sgCONTROL nuclear extracts plotted on (top) log scale or (bottom) linear scale.

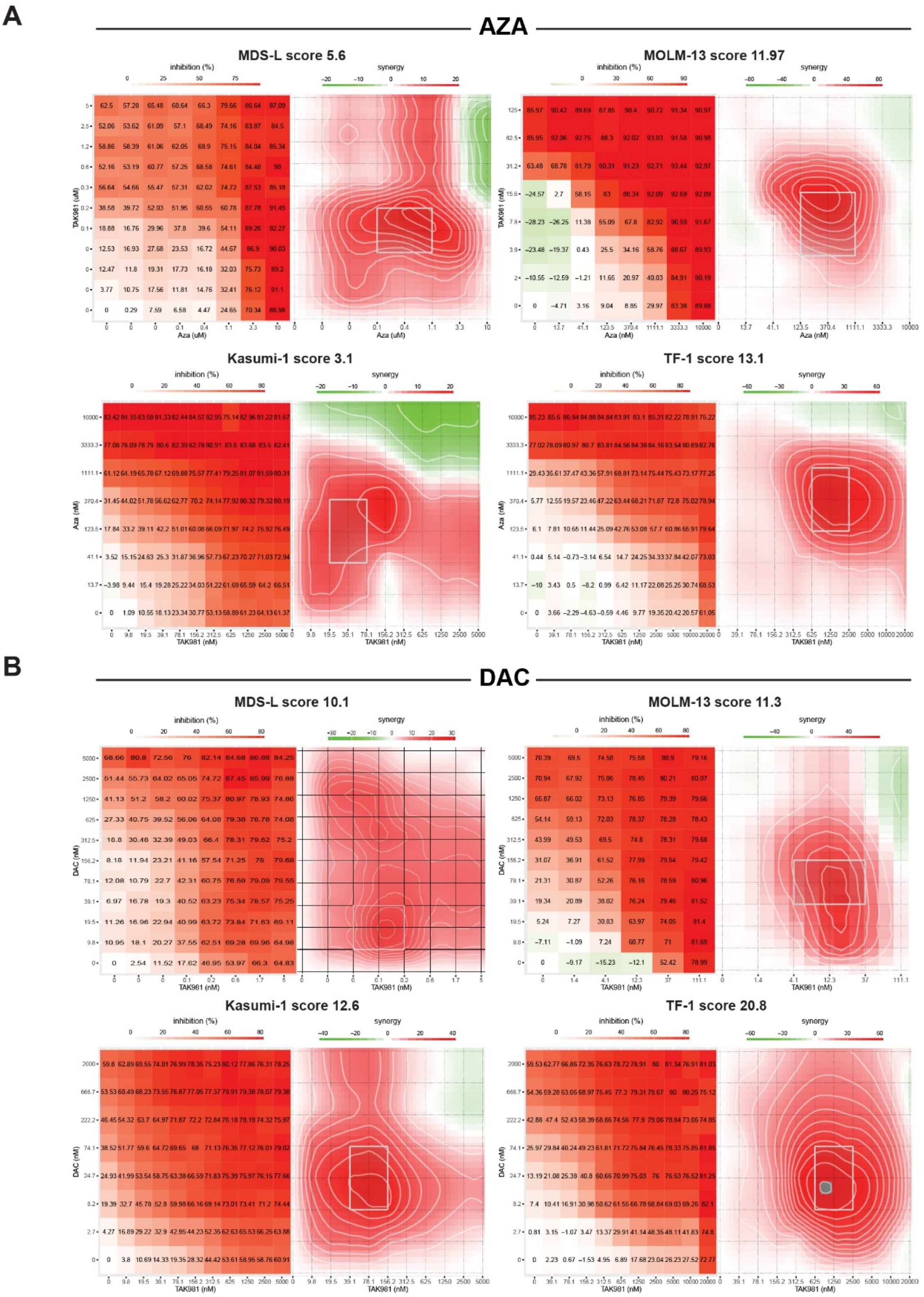
A
AZA
MDS-L score 5.6
MOLM-13 score 11.97
Kasumi-1 score 3.1
TF-1 score 13.1
inhibition (%)
synergy
TAK981 (uM)
Aza (uM)
TAK981 (nM)
Aza (nM)
B
DAC
MDS-L score 10.1
MOLM-13 score 11.3
Kasumi-1 score 12.6
TF-1 score 20.8
DAC (nM)

Figure S7

**Figure S7. SUMOylation blockade synergizes with HMAs in MDS and AML.**

Normalised dose survival matrices (left) and Synergy distribution plots (right) generated by SynergyFinder from 96-well cultures in which cross-diluted TAK-981 and (A) AZA or (B) DAC was added daily on days 1–4, and cell survival was measured by MTS assay on day 5.

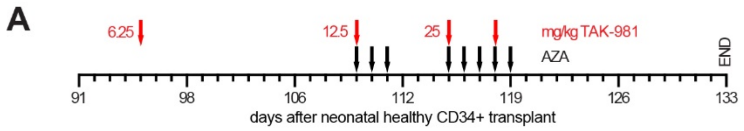
A
6.25
12.5
25
mg/kg TAK-981
AZA
END
91
98
106
112
119
126
133
days after neonatal healthy CD34+ transplant

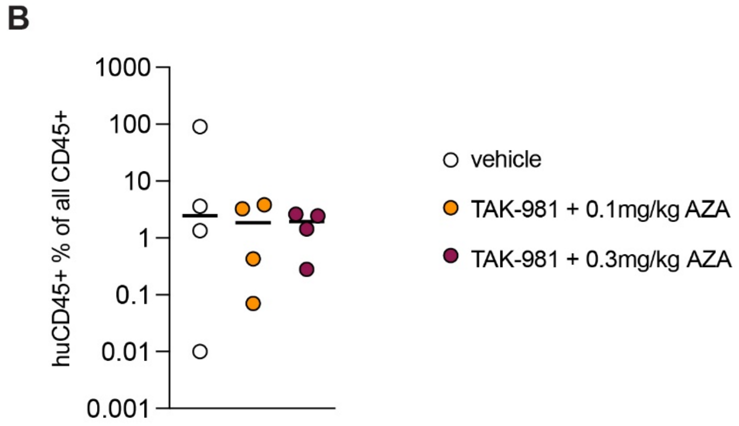
B
1000
100
10
1
0.1
0.01
0.001
huCD45+ % of all CD45+
vehicle
TAK-981 + 0.1mg/kg AZA
TAK-981 + 0.3mg/kg AZA

**Figure S8**

**Figure S8. Human bone marrow cells engrafted in MISTRG mice persist following SUMOylation blockade combined with AZA.**

MISTRG neonates were engrafted with 2,000 cord blood CD34+ cells, randomised between treatment groups based on sex and huCD45+ blood engraftment, then treated with drugs as indicated. (A) Experimental timeline. (B) Frequencies of huCD45+moCD45- cells amongst all CD45+ cells in MISTRG bone marrow after exposure to TAK-981 i.p. plus AZA s.c. *in vivo*.
